# Real time visualisation of conjugation reveals the molecular strategy evolved by the conjugative F plasmid to ensure the sequential production of plasmid factors during establishment in the new host cell

**DOI:** 10.1101/2022.09.06.506729

**Authors:** Agathe Couturier, Chloé Virolle, Kelly Goldlust, Annick Berne-Dedieu, Audrey Reuter, Sophie Nolivos, Yoshiharu Yamaichi, Sarah Bigot, Christian Lesterlin

## Abstract

DNA conjugation is a contact-dependent horizontal gene transfer mechanism responsible for disseminating drug resistance among bacterial species. Conjugation remains poorly characterised at the cellular scale, particularly regarding the reactions occurring after the plasmid enters the new host cell. Here, we use live-cell microscopy to visualise the intracellular dynamics of conjugation in real time. We reveal that the transfer of the plasmid in single-stranded DNA (ssDNA) form followed by its conversion into double-stranded DNA (dsDNA) are fast and efficient processes that occur with specific timing and subcellular localisation. Notably, the ss-to-dsDNA conversion is the critical step that governs the timing of plasmid-encoded protein production. The leading region that first enters the recipient cell carries single-stranded promoters that allow the early and transient synthesis of leading proteins immediately upon entry of the ssDNA plasmid. The subsequent ss-to-dsDNA conversion turns off leading gene expression and licences the expression of the other plasmid genes under the control of conventional double-stranded promoters. This elegant molecular strategy evolved by the conjugative plasmid allows for the timely production of factors sequentially involved in establishing, maintaining and disseminating the plasmid.

## Introduction

Bacterial DNA conjugation is a widespread horizontal gene transfer mechanism in which genetic information is transmitted from a donor to a recipient cell by direct contact (Cruz et al., 2010; Grohmann et al., 2003; Lederberg and Tatum, 1946; Virolle et al., 2020). Conjugation is responsible for the intra- and inter-species dissemination of various metabolic properties and accounts for 80% of acquired resistances in bacteria (Barlow, 2009). The F plasmid was the first conjugative element discovered (Lederberg and Tatum, 1946; Tatum and Lederberg, 1947) and is now documented as the paradigmatic representative of a large group of conjugative plasmids widespread in *Escherichia coli* and other Enterobacteriaceae species, in which they are associated with the dissemination of colicins, virulence factors, and antibiotic resistance (Fernandez-Lopez et al., 2016; Johnson et al., 2016; Lanza et al., 2014). Due to their fundamental and clinical importance, F-like plasmids have been the focus of extensive studies that provided a detailed understanding of the molecular reactions and factors involved in their transfer by conjugation (*see* (Cruz et al., 2010; Virolle et al., 2020).

Within the donor cell, the relaxosome components, including the integration host factor IHF, plasmid-encoded accessory proteins TraY, TraM and the multifunctional relaxase TraI (VirD2), are recruited to the origin of transfer (*oriT*) of the F plasmid (Howard et al., 1995; Nelson et al., 1993; Schildbach et al., 1998). The relaxosome complex is then recruited to the Type IV secretion system (T4SS) by the coupling protein TraD (VirD4), resulting in the formation of the pre-initiation complex (Beranek et al., 2004; Gomis-Rüth et al., 2004; Lang and Zechner, 2012; Llosa et al., 2003; Schröder and Lanka, 2005). It is proposed that the establishment of the mating pair induces a still uncharacterised signal that activates the pre-initiation complex. Then, TraI introduces a site- and strand-specific DNA cut (nick) into the plasmid’s *oriT* and remains covalently bound to the 5’ phosphate end. TraI also serves as a helicase that extrudes the ssDNA plasmid to be transferred, called the T-strand (Clewell and Helinski, 1970; Dostál and Schildbach, 2010; Everett and Willetts, 1980; Lanka and Wilkins, 1995; Matson and Morton, 1991; Matson and Ragonese, 2005; Reygers et al., 1991; Traxler and Minkley, 1988; Willetts and Skurray, 1980). It was initially suggested and later confirmed that two relaxases are required to carry out these functions (Dostál et al., 2011; Ilangovan et al., 2017). At this stage, the 3’OH of the T-strand serves to initiate the rolling-circle replication (RCR) that converts the intact circular ssDNA plasmid into dsDNA in the donor cell (Cruz et al., 2010; Llosa et al., 2002; Wawrzyniak et al., 2017), while the 5’phosphate bound to TraI is transferred into the recipient cell through the T4SS machinery. If the molecular structure of the T4SS has been well characterised (Christie et al., 2014; Fronzes et al., 2009; Grohmann et al., 2018; Macé et al., 2022), the way the T-strand-TraI nucleoprotein complex is translocated through the membrane of the donor and recipient cells’ membranes remain unclear.

The first transferred segment is the ∼13.5 knt leading region, carrying genes which encode the Ssb^F^ protein homolog to the chromosomally encoded essential single-strand-binding protein Ssb, the PsiB protein (<u>P</u>lasmid <u>S</u>OS <u>I</u>nhibition) (Althorpe et al., 1999a; Bagdasarian et al., 1992; Bailone et al., 1988; Dutreix et al., 1988) that inhibits SOS induction during conjugation (Baharoglu and Mazel, 2014; Baharoglu et al., 2010), and others proteins of unknown function. Remarkably, the leading region is conserved in various enterobacterial plasmids belonging to a variety of incompatibility groups (Cox and Schildbach, 2017; Golub and Low, 1985, 1986a; Golub et al., 1988; Loh et al., 1989, 1990). The adjacent and next transferred ∼17 knt maintenance region carries the ParABS-like plasmid partition system (SopABC) and the origins of vegetative replication (Bouet and Funnell, 2019; Keasling et al., 1992; Kline, 1985; Thomas, 2000). The last transferred segment of the F plasmid is the large ∼33.3 knt *tra* region that encodes all the protein factors required for plasmid DNA processing and transfer, including the relaxosome, the T4SS and the exclusion system against self-transfer (Virolle et al., 2020). Besides, F plasmids often carry cargo genes involved in various metabolic functions commonly integrated between the maintenance and the *tra* regions (Johnson et al., 2016; Lanza et al., 2014).

Once both 5’ and the 3’ ends of the T-strand have been internalised into the recipient cell, now called a transconjugant, the ssDNA plasmid is circularised by TraI and subsequently converted into dsDNA by the complementary strand synthesis reaction (Chandler et al., 2013; Dostál and Schildbach, 2010; Dostál et al., 2011; Draper et al., 2005; Garcillán-Barcia et al., 2007). The ss-to-dsDNA conversion reaction is required for plasmid replication and partition and is, therefore, critical to plasmid stability in the new host cell lineage.

The above-described mechanistic model is well-documented; however, the real time dynamics and intracellular organisation of conjugation remain largely undescribed in the live bacterium. In particular, we know very little about the subcellular localisation and timing of the reactions in the recipient cell, including the ssDNA plasmid entry, the ss-to-dsDNA conversion and plasmid gene expression. Regarding the latter, early works reported that some leading genes (*ssb*^*F*^ and *psiB* in F plasmid, and *ssb*^*ColIb-P9*^, *psiB* and *ardA* in ColIb-P9 plasmid) are expressed rapidly after entry of the plasmid in the acceptor cell (Althorpe et al., 1999b; Bagdasarian et al., 1992; Cram et al., 1984; Dutreix et al., 1988; Golub and Low, 1986a; Jones et al., 1992). *In vitro* work by Masai *et al*. (Masai and Arai, 1997) showed that the single-stranded form of the non-coding F*rpo* sequence, located in the F plasmid leading region, folds into a stem-loop structure that reconstitutes canonical -10 and -35 boxes. This promoter sequence can recruit the *E. coli* RNA polymerase that initiates RNA synthesis in *in vitro* assays (Masai and Arai, 1997). Sequences homologous to F*rpo* were also found in the leading region of ColIb-P9 (Bates et al., 1999; Nasim et al., 2004). These observations led to the proposal that F*rpo*-like sequences could act as ssDNA promoters initiating the early transcription of leading genes when the plasmid is still in ssDNA form. Whether this regulation mechanism happens during *in vivo* conjugation remains to be demonstrated.

In this study, we use live-cell microscopy imaging to visualise the complete transfer sequence of the native F plasmid between *E. coli* K12 strains. We inspect the key steps of conjugation using specifically developed genetic reporters, including a fluorescent fusion of the chromosomally encoded single-strand-binding protein Ssb (Ssb-Ypet) to monitor the ssDNA transfer, the mCherry-ParB/*parS* system to reveal the ss-to-dsDNA conversion and subsequent plasmid duplication, and translational fluorescent fusions to quantify and time plasmid-encoded production in the new host cell (Goldlust et al., 2022; Nolivos et al., 2019). This approach uncovers the choreography of conjugation reactions in live bacteria and provides new insights into the interplay between plasmid processing and gene expression.

## Results

### Dynamics of the ssDNA plasmid during transfer

We monitored the dynamic localisation of a fluorescent fusion of the chromosomally encoded single-strand-binding protein Ssb (Ssb-Ypet) in donor and recipient cells, during vegetative growth and conjugation (Figure 1A-B and Figure S1). During vegetative growth, Ssb-Ypet forms discrete foci at midcell and quarter positions within the inner region of donors and recipient cells (Figure 1C and Figure S2A-B). These Ssb foci, termed Ssb replicative foci hereafter, are associated with the ssDNA that follows the replication forks onto the nucleoid DNA (Reyes-Lamothe et al., 2008, 2010). During conjugation, the intracellular localisation of Ssb changes dramatically. As previously reported (Goldlust et al., 2022; Nolivos et al., 2019), the entry of the ssDNA plasmid in the recipient cell, now called a transconjugant, triggers the recruitment of Ssb molecules and the formation of bright membrane-proximal foci, we termed Ssb conjugative foci (Figure 1B, Figure S1). Here, we also observe the formation of Ssb conjugative foci in the donor cells, thus revealing the presence of ssDNA plasmid on each side of the conjugation pore during transfer (Figure 1B, Figure S1). Foci localisation analysis reveals that plasmid exit and entry occur at specific membrane positions within the mating pair cells. Ssb conjugative foci are mainly distributed along the donor cells’ side with a noticeable enrichment at the cell quarter positions (Figure 1C, Figure S2A-B), reflecting the preferred position for the exit of the ssDNA plasmid through active conjugation pores. By contrast, ssDNA plasmid entry predominantly occurs within the polar regions of the transconjugant cells (Figure 1C, Figure S2A-B). Our data also allow us to address whether conjugation occurs at a specific cell cycle stage. Analysis of cell length as a proxy of cell age reveals that donor and recipient cells engaged in plasmid transfer exhibit similar length distribution than during vegetative growth (Figure 1D). This shows that conjugation is cell-cycle independent as the donors can give, and recipients can acquire the plasmid at any stage of their cell cycle, from birth to cell division.

**Figure 1.**
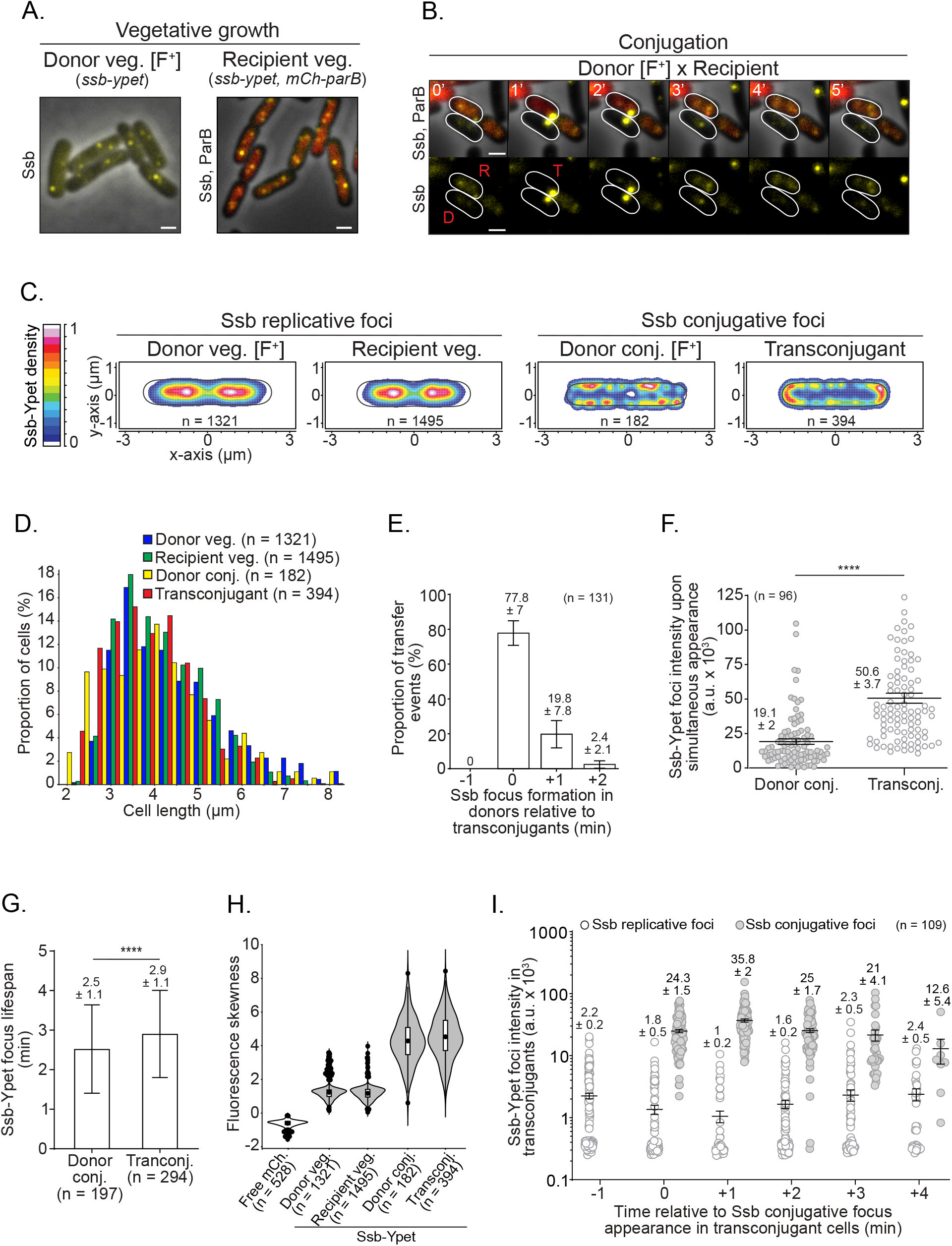
Real time dynamics of ssDNA plasmid transfer from donor to recipient cells. (**A**) Snapshot microscopy imaging of donor and recipient strains carrying the endogenous *ssb-ypet* fusion gene on the chromosome during vegetative growth. The recipient cells also produce the mCh-ParB fluorescent protein from the pSN70 plasmid that diffuses freely into the cytoplasm in the absence of the F plasmid carrying the *parS-*binding site. Scale bars 1μm. (**B**) Time-lapse microscopy images of conjugation performed in microfluidic chamber showing a plasmid transfer event between a donor (D) and a recipient cell (R) that is converted into a transconjugant (T). The ssDNA plasmid transfer is reported by the formation of paired bright membrane-associated Ssb-Ypet foci in both donor and tranconjugant cells. Scale bars 1μm. Additional transfer events are presented in Figure S1. (**C**) 2D localisation heatmaps of Ssb-Ypet fluorescent protein in donor, recipient cells in vegetative growth and in donor and transconjugant cells during conjugation. Heatmaps correspond to the merge and normalisation by the cell length of (n) individual cells from at least three biological replicates. The density scale bar is shown on the left. (**D**) Cell length distribution histogram of donor and recipient cells during vegetative growth, and of donor and transconjugant cells during conjugation (n cells analysed from at least three independent experiments). (**E**) Apparition timing of the Ssb conjugative focus in donor relative to transconjugant cells. Histograms represent the proportion of individual transfer events in which the Ssb focus appears in the donors before (−1 min), at the same time of (0 min) or after (+1 min; +2 min) the formation of a Ssb focus in the transconjugants. The number (n) of individual transfer events analysed from three independent experiments is indicated (**F**) Jitter plot of the fluorescence intensity of Ssb-Ypet conjugative foci upon simultaneous formation in donor and transconjugant cells. The number of foci analysed from three independent experiments (n) is indicated. *P-value* significance from Mann-Whitney statistical test is indicated by ****(P≤ 0.0001). (**G**) Histograms of Ssb-Ypet conjugative foci lifespan in donor and transconjugant cells measured at the single-cell level. *P-value* significance from Mann-Whitney statistical test is indicated by ****(P = 0.0001). The number (n) of cells analysed from at least five independent experiments is indicated. (**H**) Violin plots showing the fluorescence skewness of a free mCherry produced from a plasmid and of the chromosomally encoded Ssb-Ypet in donor and recipient cells during vegetative growth or donor and transconjugant cells during conjugation. The median, quartile 1 and quartile 3 are indicated by horizontal lines and the mean by a black dot. Black dots above and below the max and min values correspond to outlier cells. The number of cells analysed (n) from one representative experiment is indicated. (**I**) Jitter plot showing the evolution of the intensity of Ssb-Ypet replicative and conjugative foci in transconjugant cells in the course of the conjugation process. Time 0 minute corresponds to the appearance of the Ssb-Ypet conjugative focus in recipient cells. The number of cells analysed (n) from three independent experiments is indicated. Donor (LY1007), recipient (LY358), transconjugant (LY358 after F*wt* acquisition from LY1007); free mCherry producing strain (LY318).

In 77.8 ± 7 % (n = 131) of individual plasmid transfer events visualised by time-lapse imaging (1 min/frame), Ssb conjugative foci appear in the donor and transconjugant cells on the same frame (Figure 1E). In these cases, Ssb conjugative foci are, on average brighter in the transconjugant than in the donor cells, reflecting the relative amount of ssDNA plasmid on each side of the conjugation pore (Figure 1F). In the remaining 22.2 % of transfer events, Ssb conjugative foci first appear in the transconjugant and then in the donor one or two minutes later (Figure 1E). The delayed accumulation of ssDNA in the donor relative to the recipient is corroborated by the quantification of a 2.9 ± 1.1 min (n = 294) average lifespan of Ssb-Ypet conjugative foci in the transconjugants, compared to 2.5 ± 1.1 min (n = 197) in the donor cells (Figure 1G). These data indicate that the appearance of conjugative foci is asynchronous in the mating pair cells and suggest a specific sequence of ssDNA transfer. The first segment of the T-strand generated by the helicase activity of TraI in the donor cell does not dwell long enough to recruit Ssb molecules and is immediately transferred to the recipient. Only after this brief transfer stage does the ssDNA accumulates on the donor’s side as well, where it can correspond to either or both the non-transferred plasmid strand or to the T-strand. This implies that the rate of ssDNA formation by TraI helicase activity is faster than that of ssDNA removal by the RCR and transfer through the T4SS (See discussion).

The internalisation of a large amount of ssDNA plasmid provokes the massive recruitment of the intracellular pool of Ssb molecules at the periphery of the donor and transconjugant cells. This change in Ssb-Ypet subcellular distribution is revealed by skewness analysis, which provides a non-biased measure of the asymmetry of fluorescence distribution within the cells without a requirement for threshold-based foci detection (Figure 1H). Cells producing a free mCherry (mCh) exhibit a low skewness corresponding to the homogeneous pixel fluorescence distribution inside the cell’s cytoplasm. During vegetative growth, Ssb-Ypet fluorescence is partly diffuse in the cytoplasm and partly locally concentrated within replicative foci, resulting in a skewness of ∼1.2. By comparison, Ssb-Ypet exhibits a strong skewness of ∼4.1 in donors and transconjugants during plasmid transfer, reflecting the increased proportion of Ssb molecules clustered within foci. Hence, we wondered what part of Ssb molecules are contained within conjugative foci and if their formation was associated with a depletion of Ssb within replicative foci in the transconjugant cell. To address this question, we performed Ssb-Ypet foci automatic detection and brightness quantification during plasmid transfer (Figure 1I). We observe that one minute after the beginning of plasmid entry Ssb-Ypet replicative foci are still present but exhibit half their initial intensity, while conjugative foci are 35 times brighter. Since the total Ssb-Ypet intracellular fluorescence is unchanged during the transfer (Figure S2C), these variations can be attributable to the displacement of Ssb-Ypet molecules onto the incoming ssDNA plasmid rather than Ssb-Ypet *de novo* synthesis. This dynamic reflects that the incoming ssDNA plasmid recruits most Ssb-Ypet molecules in the acceptor cell during transfer.

It has been estimated that Ssb is present at about ∼1320 ± 420 monomers per *E. coli* cell and that a dimer of tetramers covers about 170 nt *in vivo* (Reyes-Lamothe et al., 2010). Consequently, there are not enough Ssb copies per cell to accommodate the 108 000 nucleotides ssDNA F plasmid, plus the few hundreds of nucleotides of ssDNA associated with replication forks (∼650 nt at 22ºC (Lohman and Ferrari, 1994)). This raises the possibility that the reduced availability of Ssb molecules during plasmid entry could provoke a transitory disturbance of the host chromosome DNA replication. One way to address this question *in vivo* is to monitor a fluorescent fusion of the *β*_*2*_-clamp replisome component (mCh-DnaN), which is diffuse in the cytoplasm of non-replicating cells and forms discrete replisome-associated foci during DNA replication progression (Moolman et al., 2014; Reyes-Lamothe et al., 2008, 2010). Microscopy imaging and skewness analysis showed no change in DnaN localisation pattern before, during or after Ssb conjugative foci formation (Figure S2D). This indicates that Ssb recruitment onto the incoming ssDNA plasmid does not result in the collapse of the replication fork. Whether the rate of DNA replication is affected during this transient and short process remains a possibility.

### ss-to-dsDNA conversion and subsequent plasmid replication in the transconjugant cells

The conversion of the newly acquired ssDNA plasmid into dsDNA by the complementary strand synthesis reaction and the subsequent plasmid duplication events were analysed using the *parS/*ParB DNA labelling system (Goldlust et al., 2022; Nolivos et al., 2019). The *parS* binding site is inserted in the F plasmid, while the ParB binding protein fluorescently labelled with the mCherry (mCh-ParB) is produced from a plasmid in recipient cells only. Under the microscope, the ss-to-dsDNA conversion is reported by the disappearance of the Ssb-Ypet conjugative focus and the formation of a mCh-ParB focus in the transconjugant cells (Figure 2A). We first performed time-lapse imaging (1 min/frame) to visualise the success rate and timing of ss-to-dsDNA conversion after ssDNA entry (Figure 2B). Analysis shows that the appearance of the Ssb-Ypet conjugative focus is followed by the formation of the mCh-ParB focus in 83.3 ± 2.3 % (n = 311) individual transconjugant cells analysed, indicating that the vast majority of internalised ssDNA plasmids are successfully converted into dsDNA plasmids (Figure 2C). Notably, we observe that 40 ± 3.2 % (n = 286) of transconjugant cells where the newly acquired ssDNA plasmid has already been converted into dsDNA subsequently receive additional ssDNA (Figure 2D, Figure S3A). We quantify that 92 ± 3.1 % of these multiple ssDNA acquisition events originate from the same donor, among which 79 ± 5.3 % appear to take place at the same membrane position, suggesting that they occur through the same conjugation pore (Figure S3A). The evidence for multiple transfers within an established mating pair demonstrates that a single donor can successively give several copies of the T-strand and that transconjugants in which the ss-to-dsDNA conversion has already been achieved do not become instantly refractory to *de novo* plasmid acquisition. Accordingly, establishing immunity to conjugation by transconjugant cells is expected to require the production of the plasmid-encoded exclusion proteins TraS and TraT.

**Figure 2.**
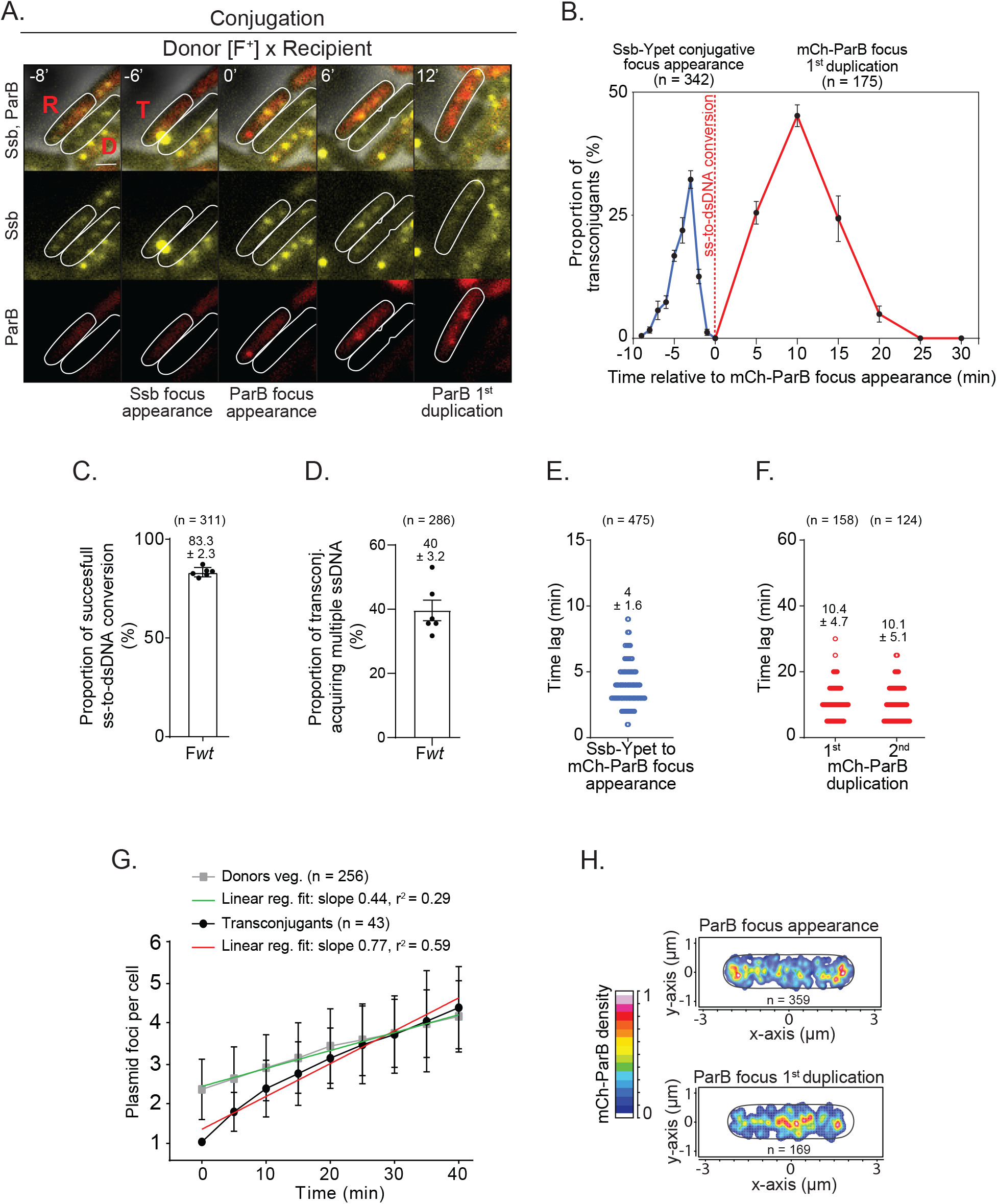
Timing and spatial localisation of the ss-to-dsDNA conversion and plasmid duplication in transconjugant cells. **(A)** Time-lapse microscopy images performed in microfluidic chamber showing the transfer of the ssDNA plasmid reported by the formation of the Ssb-Ypet conjugative foci in both donor (D) and recipient (R) cells, followed by the ss-to-dsDNA conversion reflected by the appearance of a mCh-ParB focus in transconjugant (T) cells. Scale bar 1μm. (**B**) Single-cell time-lapse quantification of Ssb-Ypet focus appearance (blue line) and mCh-ParB focus first duplication (red line) with respect to the ss-to-dsDNA conversion revealed by mCh-ParB focus formation in transconjugant cells (0 min). The number of conjugation events analysed (n) from seven independent experiments is indicated. (**C**) Histogram showing the frequency of successful ss-to-dsDNA conversion reflected by the conversion of the Ssb-Ypet conjugative foci into a mCh-ParB focus. The mean and SD are calculated from (n) individual transfer events from six biological replicates (black dots). (**D**) Histogram showing the percentage of transconjugant cells with a mCh-ParB focus that acquire multiple ssDNA plasmids as revealed by the successive appearance of an additional Ssb-Ypet conjugative focus. The mean and SD are calculated from (n) individual transconjugant cells from six biological replicates (black dots). (**E**) Scatter plot showing the time lag between the appearance of the Ssb-Ypet focus and the mCh-ParB focus in transconjugant cells. The mean and SD calculated from (n) individual ss-to-dsDNA conversion event (blue circles) from seven biological replicates are indicated. (**F**) Scatter plot showing the time-lag between the apparition of the mCh-ParB focus and its visual duplication in two foci (1^st^ duplication), and in three or four foci (2^nd^ duplication). The mean and SD calculated from (n) individual duplication events (red circles) from at least six biological replicates are indicated. (**G**) Single-cell time-lapse quantification of the number of F foci per cell in F-carrying donor strain during vegetative growth and in transconjugants after F plasmid acquisition. For donor, the number of F foci per cell (reflected by the number of SopB-sfGFP foci) with respect to cells birth (t = 0 min) is shown (grey curve). For transconjugants the number of F foci per cell (reflected by the number of mCh-ParB foci) with respect to mCh-ParB focus appearance (t = 0 min) is shown (black curve). Mean and SD calculated from (n) individual cells from four biological replicates are indicated, together with curves’ linear fitting lines for donors (green) and transconjugants (red). F-carrying donor strain (LY834), Transconjugant (LY358 after F*wt* acquisition). (**H**) 2D localisation heatmaps of the mCh-ParB focus at the time of its appearance (top) and just before its duplication into two foci (bottom). Heatmaps correspond to the merge and normalisation by the cell length of (n) individual transconjugant cells from seven biological replicates. (**A-F and H**) F*wt* donor (LY1007), recipient (LY358), transconjugant (LY358 after F*wt* acquisition).

Considering successful ss-to-dsDNA events only, we calculate an average 4 ± 1.6 min (n = 475) time lag between the appearance of the Ssb-Ypet conjugative focus and the formation of the mCh-ParB focus (Figure 2E). This period reflects the time required for the completion of a reaction cascade that comprises the complete internalisation of the ssDNA plasmid, the circularisation of the ssDNA plasmid by TraI, the initiation and completion of the complementary strand synthesis replication, and the recruitment of ParB molecules on the *parS* site in dsDNA form. Though our system does not allow evaluating each step’s contribution, results show that the complete sequence of reactions is achieved within a relatively short and consistent period.

Next, we first performed time-lapse imaging (5 min/frame) to examine the timing of plasmid duplication in transconjugant cells (*i*.*e*., replication and visual separation of the plasmid copies) (Figure 2B). We estimate an average of 10.4 ± 4.7 min (n = 158) period between the ssDNA-to-dsDNA conversion and the first plasmid duplication event (from one to two mCh-ParB foci) and similar 10.1 ± 5.1 min (n = 124) between the first and the second duplication event (from two to three or four mCh-ParB foci) (Figure 2F). We then decided to compare the rate of plasmid duplication in transconjugants to the rate of plasmid duplication in a vegetatively growing F-carrying donor strain. To do so, we plotted the number of plasmid foci per cell from the ss-to-dsDNA conversion (mCh focus appearance) to cell division in transconjugants and from cell birth to cell division in F-carrying donor cells (Figure 2G). Results show that the number of F per cell increases significantly faster in transconjugant cells than in vegetatively growing F-carrying cells (75 % increase of the fit curve slope), yet to reach a similar final number of ∼4 ± 1 copies per cell before division (Figure 2G). F copy number, like chromosome replication, is known to be controlled by the cell cycle progression, where initiation occurs when a constant mass per origin is achieved (Keasling et al., 1991). Therefore, our observations are consistent with the interpretation that when a single plasmid copy arrives in a recipient cell that can be at any cell cycle stage, plasmid replication initiation is unrepressed until the specific number of plasmid copies per cell mass is restored. This accelerated plasmid replication allows for the rapid increase in F copy number before the division of the transconjugant cells, thus facilitating the segregation of plasmid copies to daughter cells.

Localisation analysis reveals that the ss-to-dsDNA conversion and the first duplication event occur at distinct subcellular positions. The initial mCh-ParB focus preferentially appears in the polar region of the transconjugant cell, comparable to the ssDNA’s entry location (compare Figure 2H to Figure 1C and Figure S3B to Figure S2A). A noticeable difference is that mCh-ParB foci appear less peripheral, indicating that they are not as close to the cell membrane as Ssb-Ypet conjugation foci (compare Figure 2H to Figure 1C, and Figure S3C to Figure S2B). We observe that the mCh-ParB focus subsequently migrates to the midcell position before duplication (Figure 2H, Figure S3B-C). These data show that the two DNA synthesis reactions involved in plasmid processing (*i*.*e*., ss-to-dsDNA conversion and plasmid replication) are separated in time and space in the new host cell. The recruitment of the complementary strand synthesis machinery and the ss-to-dsDNA replication reaction occur in the vicinity of the polar position of entry of the ssDNA plasmid, while plasmid replication occurs in the midcell region. Altogether, these analyses reveal that plasmid processing steps (ssDNA entry, ss-to-dsDNA conversion and plasmid replication) occur at specific intracellular positions within the new host cell and follow a precise chronology.

### Program of plasmid-encoded protein production in transconjugant cells

We constructed *superfolder gfp* (*sfgfp*) C-terminal translational fusions to several genes located in the different functional regions of the F plasmid to examine the production timing of plasmid-encoded proteins in transconjugant cells, which we use to get insights into the timing of plasmid gene expression (Figure 3A, Figure S4A). *YgfA, ygeA, psiB, yfjB, yfjA* and *ssb*^*F*^ are located in the leading region and are transferred in order after the origin of transfer *oriT*. The *sopB* gene is part of the SopABC partition system and is located in the maintenance region. The *traM, traC, traS* and *traT* genes are located in the *tra* region that encodes factors involved in plasmid transfer. TraM is the accessory protein of the relaxosome complex that is recruited to the *oriT* (Di Laurenzio et al., 1992); TraC is the traffic ATPase organised as a hexamer of dimers docked to the cytoplasmic faces of the T4SS (Hu et al., 2019); TraS and TraT correspond to the F plasmid exclusion (immunity) system that protects against self-transfer (Achtman et al., 1977; Jalajakumari et al., 1987; Manning et al., 1980).

**Figure 3.**
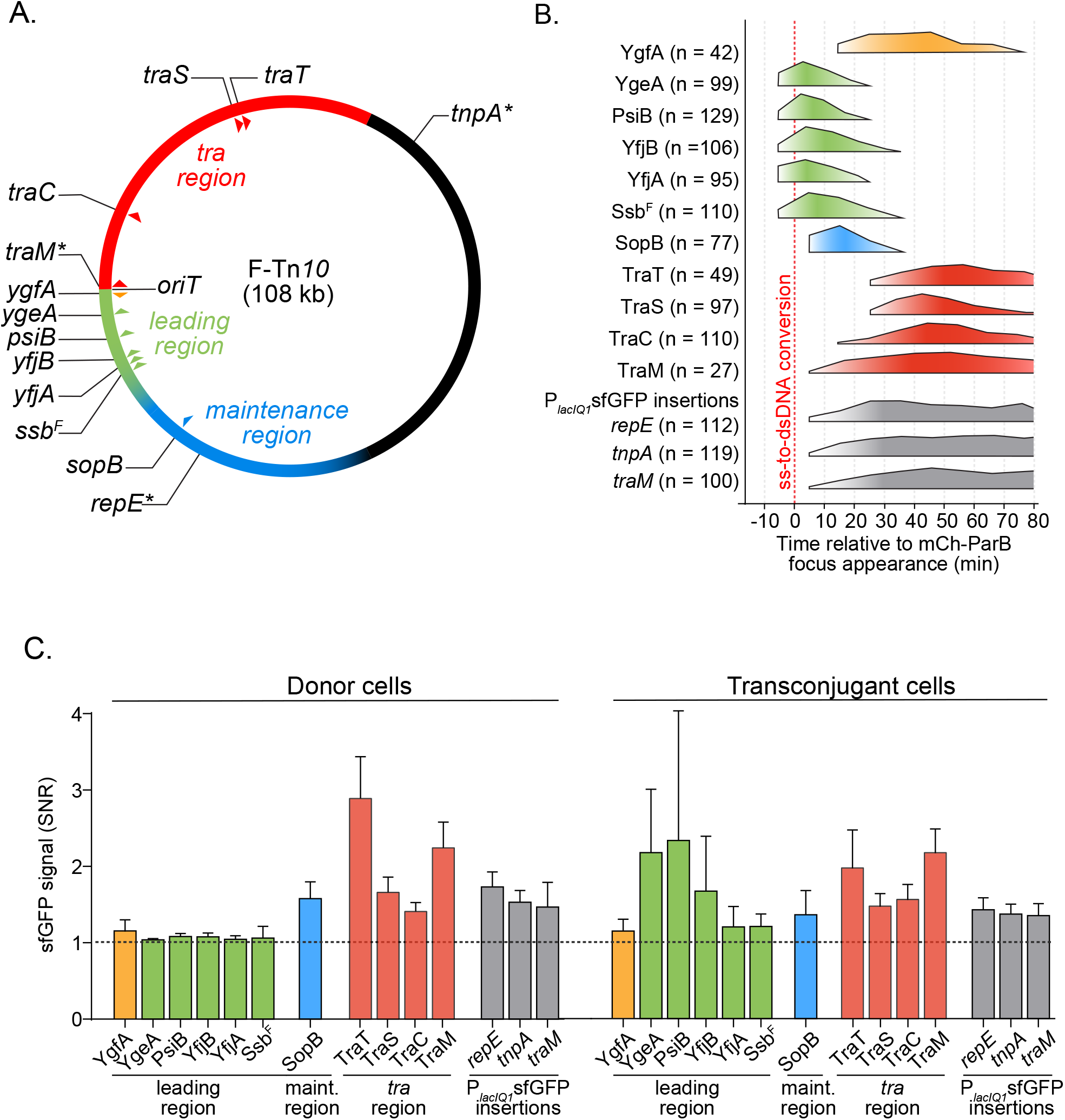
Timing of plasmid-encoded proteins production in transconjugant cells. (**A**) Genetic map of the 108 kb F plasmid indicating the leading (green), Tra (red) and maintenance (blue) regions, and the positions of the studied genes (triangles). Stars represent the genetic location of the P_*lacIQ1*_*sfgfp* insertions. (**B**) Summary diagram of the production timing of each plasmid-encoded protein fusions in transconjugant cells with respect to the timing of ss-to-dsDNA conversion reflected by mCh-ParB focus appearance (0 min). The diagram represent data from the foldchange increase in sfGFP signal from Figure S5. Orange/green, blue and red colours correspond to production of proteins from the leading, maintenance and transfer region respectively. Timings of the cytoplasmic sfGFP production from the P_*lacIQ1*_ promoter inserted in the *repE-sopA* (*repE*), *tnpA-ybaA* (*tnpA*) and *traM-traJ* (*traM*) intergenic regions are represented in grey. The number (n) of individual transconjugant cells from at least three biological replicates analysed is indicated. (**C**) Histograms showing the intracellular green fluorescence (SNR) for each sfGFP fusions and reporters within vegetatively growing donor (left) and transconjugant cells (right) at the maximum SNR value from Figure S5. Means and SD calculated from the same individual transconjugant cells as in (B) are indicated. Donors of F derivatives (see Table S1), Recipient (LY358).

We first performed time-course experiments where microscopy snapshot images of the conjugating population were acquired 1, 2, 4 and 6 hours after mixing donors and recipient cells. For each time point, the frequency of transconjugants (T/R+T) was directly measured at the single-cell level from the proportion of recipient cells exhibiting diffuse mCh-ParB fluorescence (R) or transconjugant cells harbouring mCh-ParB foci (T,), and the intracellular green fluorescence Signal to Noise Ratio (SNR) was automatically measured (Figure S4B-D). This snapshot analysis shows that all F plasmid derivatives carrying sfGFP fusions retained their transfer ability and raised frequencies of transconjugants between 57 and 93 % after 6 hours of mating. Also, fusion-carrying plasmid acquisition is systematically followed by an increase in sfGFP signal in transconjugant cells, with highly variable timing and levels (Figure S4B-D).

Better resolution of the production level and timing of sfGFP fusions with respect to the ss-to-dsDNA conversion (appearance of the mCh-ParB focus) in individual transconjugant cells was obtained using time-lapse imaging of conjugation performed in the microfluidic chamber (Movie S1 and S2). We performed transconjugant cell detection and quantification of the intracellular sfGFP SNR cells over time (Figure S5A-D). When the transconjugant cell divided, we continued fluorescence quantification in the resulting daughter cells to monitor sfGFP production over a longer period. From this raw data, we calculated the fold-increase in SNR per ten-minute interval, where a fold-increase superior to one reveals that the fusions are being produced in the transconjugants (Figure S5A-D). These data were finally translated into a comprehensive diagram presenting the production time windows for each fusion in transconjugant cells relative to the ss-to-dsDNA conversion event (Figure 3B). This analysis reveals that fusions belonging to the different plasmid regions exhibit specific production timings with respect to plasmid processing steps.

Remarkably, we detect the synchronous production of the leading YgeA, PsiB, YfjB, YfjA and Ssb^F^ fusion proteins even before the appearance of the mCh-ParB focus (Figure 3B and Figure S5A). Furthermore, the production of these fusions is only transient as it peaks at ∼5 minutes and stops 25-35 minutes after the ss-to-dsDNA conversion event. This unexpected observation indicates that leading fusions start being produced when the plasmid is still in ssDNA form and stops rapidly after the plasmid is converted into dsDNA form. An interesting exception is YgfA-sfGFP, for which production is only detected in the 10-20 minutes interval after mCh-ParB focus appearance. The *ygfA* gene is the closest to the *oriT* and is, therefore, the first gene to be transferred into the recipient (Figure 3A, Figure S4A). However, *ygfA* gene orientation is opposite to other tested leading genes, meaning that the T-strand does not correspond to the template strand for *ygfA* transcription. Consequently, and consistent with our observations, *ygfA* expression can only occur after synthesising the complementary template strand by the ss-to-dsDNA conversion.

The ss-to-dsDNA conversion is followed by the production of maintenance and Tra proteins, starting with SopB and TraM, then TraC, and eventually TraS and TraT fusions (Figure 3B, Figure S5B-C). The production of these fusions is expected to require the presence of the plasmid in dsDNA form since the corresponding genes are known to be controlled by dsDNA promoters (*P*_*sopAB*_ for *sopB, P*_*M*_ for *traM* and *P*_*Y*_ for *traC* and *traST*). However, what could explain the observed differences in the production timings? We addressed whether timing discrepancies could simply account for the fusions’ position on the genetic map of the F plasmid. This possibility was excluded by the observation that insertion of the constitutive fluorescent reporter *P*_*lacIQ1*_sfGFP (*sfgfp* gene under the control of the *P*_*lacIQ1*_ constitutive promoter) in the *repE-sopA, tnpA-ybaA* and *traM-traJ* intergenic regions resulted in similar sfGFP production timings, within the 0-10 minutes interval after the appearance of the mCh-ParB focus (Figure 3B, Figure S5D). Instead, we propose that the differential production timings of maintenance and *tra* genes reflect the activity and regulation of the promoters of the corresponding genes. The *sopAB* operon is under the control of the *P*_*sopAB*_ promoter, which is repressed by SopA binding. Therefore, the *P*_*sopAB*_ promoter is expected to be fully unrepressed and active in transconjugant cells devoid of SopA, thus allowing the rapid production of the SopAB partition complex required for plasmid stability and inheritance over cell divisions. The *traM* gene is controlled by the *P*_*M*_ promoter, which is weakly but constitutively active, even before its full activation by binding the TraY protein (Penfold et al., 1996). By contrast, the *P*_*Y*_ promoter that controls the expression of *traC, traS* and *traT* genes needs to be activated by the TraJ protein, encoded by the *traJ* gene under the control of its own promoter *P*_*J*_ and located upstream of *P*_*Y*_ (Virolle et al., 2020). The requirement for this activation cascade probably explains the delayed production of TraC, TraS and TraT. The additional delay between TraC and TraS/TraT fusions production could potentially reflect the relative distance of these genes to the *P*_*Y*_ promoter (5.9 kb for *traC* and 20.4 kb for *traST*).

Notably, the intracellular levels of Tra proteins within transconjugant cells reach a plateau between 60 to 90 minutes after the ss-to-dsDNA conversion and remain stable throughout our observations (Figure 3B, Figure S5C). This involves that at that point, transconjugant cells have produced the transfer machinery and the exclusion system and have most likely been converted into proficient plasmid donors. In support of this interpretation, TraM, TraC, TraS, TraT and SopB are detected at similar levels in vegetatively growing F-carrying donor cells (Figure 3C, Figure S4C-D and S5B-C). This is not the case for YgeA, PsiB, YfjB, YfjA, and Ssb^F^ leading proteins, which intracellular levels start decreasing 25-35 minutes after the ss-to-dsDNA conversion in the transconjugants, and which are not detected in vegetatively growing donor cells (Figure 3C, Figure S4B and S5A). These results are consistent with the interpretation that leading proteins are produced rapidly and only transiently upon entry of the ssDNA plasmid in the recipient cells and not when the plasmid is maintained in dsDNA form during vegetative replication.

### Single-stranded promoters allow the early expression of the leading genes in the transconjugant cell

Together with previous works (Althorpe et al., 1999b; Bagdasarian et al., 1992; Bates et al., 1999; Jones et al., 1992), the early and transiently expression of leading genes in transconjugant cells support the existence of specific sequences that would act as single-stranded promoters to initiate the transcription of leading genes from the internalised ssDNA plasmid. Using bioinformatics analysis, we identified a region upstream of the *ssb*^*F*^, *yfjA, yfjB, psiA* and *psiB* genes, which we named F*rpo2*, that shares 92% identity with the previously reported F*rpo* region (renamed F*rpo1*) located upstream *ygeA* and *ygeB* and previously characterised *in vitro* (Masai and Arai, 1997) (Figure 4A). DNA folding prediction using mFold (http://www.unafold.org) indicates that the single-stranded form of F*rpo2* can fold into a highly stable stem-loop structure that also carries canonical -10 and -35 boxes, similar to the F*rpo1* region (Figure S6A) (Masai and Arai, 1997). We addressed the effect of F*rpo1* or F*rpo2* deletions on the expression of the downstream genes in transconjugant cells using live-cell microscopy. Microscopy analysis of transconjugant cells receiving the F ΔF*rpo1 ygeA*-*sfgfp*, the F ΔF*rpo2 ssb*^*F*^-*sfgfp*, or the F ΔF*rpo2 yjfA*-*sfgfp* revealed no significant fold-increase in sfGFP fluorescence before or after the ss-to-dsDNA conversion in the transconjugant cells (Figure 4B).

**Figure 4.**
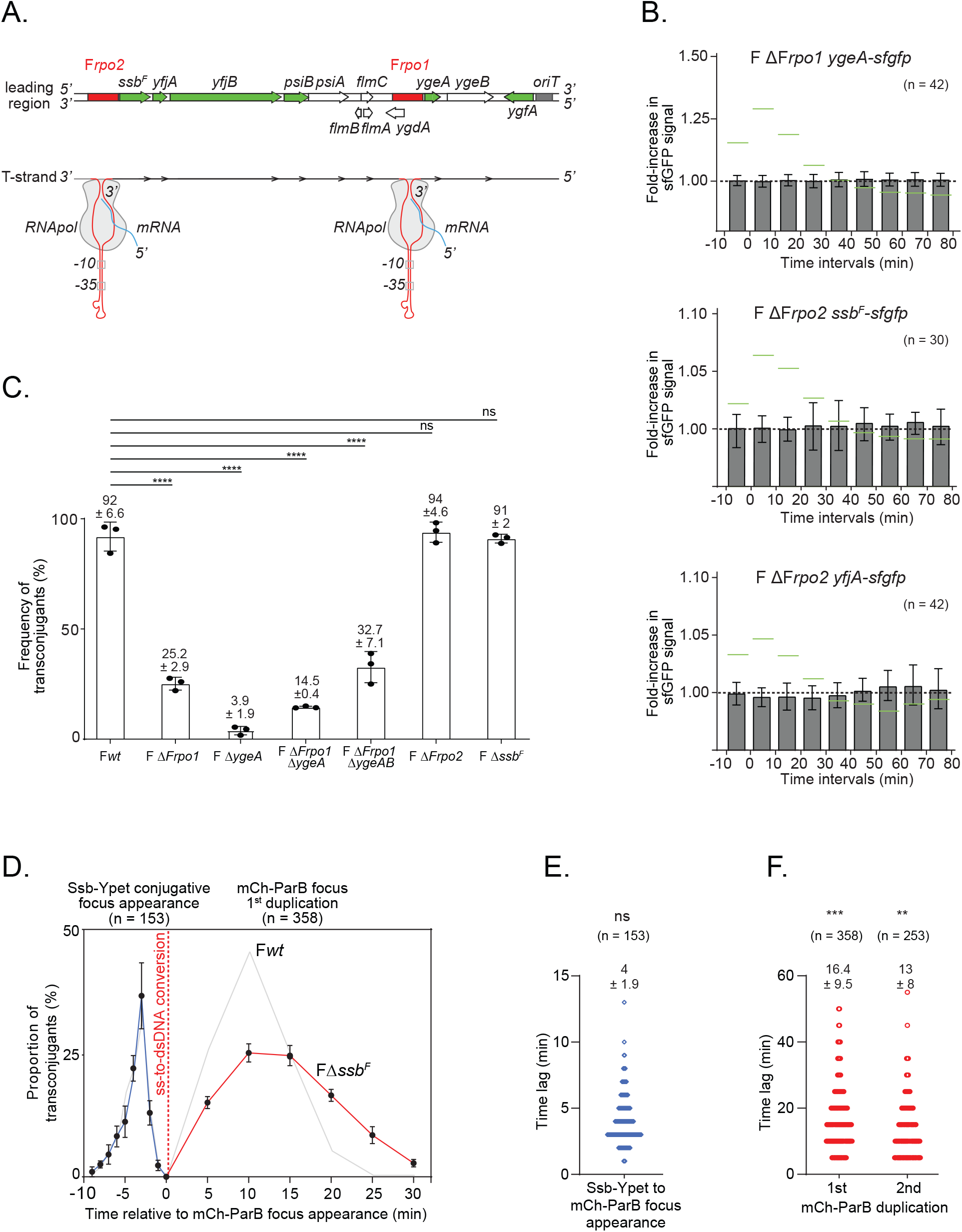
Role of leading region factors F*rpo1*, F*rpo2* and *ssb*^*F*^ in conjugation. (**A**) Genetic map of the dsDNA leading region showing the position of the genes (green for studied sfGFP fusions and white for the other genes) and F*rpo1* and F*rpo2* promoters (red) (top). The bottom diagram shows the stem-loop structure formed by the ssDNA forms of F*rpo1* and F*rpo2* promoter sequences (detailed in Figure S6). Recognition of the *-10* and *-35* boxes present in the dsDNA stem region by the RNA polymerase (RNA pol in grey) induces the initiation of transcription and the production of mRNA (blue). (**B**) Histograms of intracellular sfGFP fold increase in transconjugant after acquisition of F ΔF*rpo1 ygeA-sfgfp*, F ΔF*rpo2 ssb-sfgfp* and F ΔF*rpo2 yfjA-sfgfp*. Mean and SD are calculated from (n) individual transconjugant cells analysed from at least three independent experiments. Levels obtained with the F*wt* plasmid from Figure S5A are *wt* reported in green as a reference. Donor of F ΔF*rpo1 ygeA-sfgfp* (LY1368), F ΔF*rpo2 ssb-sfgfp* (LY1365), F ΔF*rpo2 yfjA-sfgfp* (LY1364), recipient (LY318). (**C**) Histograms of F*wt*, deletion mutants F ΔF*rpo1*, F Δ*ygeA*, F ΔF*rpo1* Δ*ygeA*, F ΔF*rpo1* Δ*ygeAB*, FΔF*rpo2* and FΔ*ssb*^*F*^ frequency of transconjugant (T/R+T) estimated by plating assays. Mean and SD are calculated from at least three independent experiments. *P*-value significance ns and \*\*\*\**P* ≤ 0.0001 were obtained from One-way ANOVA with Dunnetts multiple comparisons test. Donor of F*wt* (LY875), F ΔF*rpo1* (LY824), F Δ*ygeA* (LY160), F ΔF*rpo1* Δ*ygeA* (LY1424), F ΔF*rpo1* Δ*ygeAB* (LY1425), F ΔF*rpo2* (LY823), F Δ*ssb*^*F*^ (LY755), recipient (MS428). (**D**) Single-cell time-lapse quantification of Ssb-Ypet focus appearance (blue line) and mCh-ParB focus first duplication (red line) with respect to the ss-to-dsDNA conversion revealed by mCh-ParB focus formation in transconjugant cells (0 min) that receive the FΔ*ssb*^*F*^ plasmid. The number of conjugation events analysed (n) from five independent biological replicates is indicated. Results obtained in Figure 2B with F*wt* plasmid are reported in grey for comparison. (**E**) Scatter plot showing the time lag between the appearance of the Ssb-Ypet focus and the appearance of the mCh-ParB focus in transconjugant cells after the acquisition of the F Δ*ssb*^*F*^ plasmid. The mean and SD calculated from (n) individual ss-to-dsDNA conversion event (blue circles) from five biological replicates are indicated. P-value significance ns (>0.05 non-significant) was obtained from Mann-Whitney statistical test against results obtained with the F*wt* plasmid (Figure 2E). (**F**) Scatter plot showing the time-lag between the apparition of the mCh-ParB focus and its visual duplication in two foci (1^st^ duplication), and in three or four foci (2^nd^ duplication) in transconjugant cells after acquisition of the F Δ*ssb*^*F*^ plasmid. The mean and SD calculated from (n) individual duplication events (red circles) from eight biological replicates are indicated. P-value significance **P = 0.0023 and ***P = 0.0007 were obtained from Mann-Whitney statistical test against results obtained with the F*wt* plasmid (Figure 2F). Donor F Δ*ssb*^*F*^ (LY1068), recipient (LY358).

We then addressed the impact of F*rpo1* and F*rpo2* deletions on the efficiency of conjugation after three hours of mating, as estimated by plating assays (Figure 4C). F ΔF*rpo1* exhibits a dramatically reduced frequency of transconjugants of 25.2 ± 2.9 % compared to 92.6 ± 6.6 % for the F*wt*. Comparable results were obtained for F ΔF*rpo1* Δ*ygeAB* (32.7 ± 7.1) and F ΔF*rpo1* Δ*ygeA* (14.5 ± 0.4). Surprisingly, the single deletion of *ygeA* decreases the conjugation of efficiency even further (3.9 ± 1.9 %), and despite our multiple attempts, the deletion of *ygeB* alone could never be constructed. By contrast, the deletions of F*rpo2* or *ssb*^*F*^ have no significant impact on the conjugation efficiency. These results show that F*rpo1* and F*rpo2* are required for the early expression of the downstream genes upon plasmid entry in recipient cells during conjugation *in vivo*. However, genes under the control of F*rpo1* appear to have a more critical role in conjugation than those under the control of F*rpo2*.

### Role of the plasmid-encoded Ssb^F^ leading protein in plasmid establishment

The rapid and transient expression of leading genes upon plasmid entry strongly suggests that leading proteins have an essential role during the early steps of plasmid establishment in the new host cell. The leading region conserved in various enterobacterial plasmids encodes a homolog of the single-strand-binding protein Ssb encoded on the *E. coli* chromosome (Golub and Low, 1985, 1986b; Golub et al., 1988; Howland et al., 1989; Jones et al., 1992; Kolodkin et al., 1983). The chromosomally encoded *ssb* gene is conserved and essential in all bacterial organisms, raising the question of the *raison d’être* of plasmid-born *ssb* homologues. Early study shows that the Ssb^F^ encoded by the F plasmid can partially complement conditional mutations of the chromosomal *ssb* gene (Golub and Low, 1986b; Porter and Black, 1991). Consistently, we performed simultaneous visualisation of Ssb^F^-mCh produced from a pTrc99a-*ssb*^*F*^*-mch* plasmid and the chromosomally-encoded Ssb-Ypet (Figure S7A) and observed similar intracellular positioning (Figure S7B) confirmed by colocalisation analysis (Figure S7C). This indicates that both the plasmid Ssb^F^ and the host Ssb are recruited to the ssDNA that follows the replication forks in vegetatively growing cells. Similarly, SsbF-sfGFP also forms foci in transconjugant cells that have acquired the F *ssb*^*F*^-*sfgfp* plasmid, mainly during the first and second plasmid duplication events (Figure S7D-E). Nonetheless, the role of Ssb^F^ during conjugation is still unclear, and its deletion from the F plasmid has no significant impact on conjugation efficiency (Figure 4C).

To get further insight into the role of Ssb^F^ during conjugation, we revisited the dynamics of ssDNA entry, ss-to-dsDNA conversion and duplication of the F Δ*ssb*^*F*^ plasmid. Time-lapse microscopy image analysis reveals that Ssb^F^ deletion has no impact on the dynamics of Ssb-Ypet conjugative foci (Figure 4D) or the timing of the ss-to-dsDNA conversion (compare Figure 4E to Figure 2E). However, Ssb^F^ deletion dramatically delays the timing of plasmid duplication in transconjugant cells (compare Figure 4F to Figure 2F). The time lag between mCh-ParB appearance and the first duplication is increased by ∼58 % (from 10.4 ± 4.7 for F*wt* to 16.4 ± 9.5 for F Δ*ssb*^*F*^), and the time between the first and second plasmid replication event is increased by ∼29 % (from 10.1 ± 4.7 for F*wt* to 13 ± 8 for F Δ*ssb*^*F*^). This indicates that Ssb^F^ has a role in facilitating the first rounds of plasmid duplication in the new transconjugant cell, possibly by increasing the cellular pool of single-strand binding protein available for DNA replication. This function appears dispensable since the absence of Ssb^F^ delays plasmid duplication but does not affect the final efficiency of conjugation, at least when conjugation is performed in optimal conditions between *E. coli* MG1655 strains.

## Discussion

Our current knowledge of conjugation mainly emerges from experimental genetic, biochemical and structural studies that provided a well-documented understanding of the molecular reactions and factors involved in DNA transfer, while genomic and computational studies uncovered the diversity of conjugative plasmids and their importance in the epidemiology of antibiotics resistance dissemination. It is only recently that the application of optical microscopy has started to provide insights into the organisation of conjugation at the cellular scale (Aguilar et al., 2011; Babic et al., 2011; Babić et al., 2008; Carranza et al., 2021; Clarke et al., 2008; Goldlust et al., 2022; Lawley et al., 2002; Low et al., 2022; Nolivos et al., 2019). In this study, live-cell microscopy combined with specifically developed fluorescent reporters offers a unique view of the cellular dynamics of conjugation while providing insights into the timing and localisation of each key step.

We report the presence of ssDNA plasmid on both the donor and the recipient’s side during plasmid transfer. Noticeably, the ssDNA plasmid is not randomly positioned but instead allocated to specific subcellular locations within the mating pair cells. The exit point of the ssDNA is preferentially located on the side of the donor cell and enriched at quarter positions. This unlikely reflects a specific positioning of the T4SS machinery, which was reported to be homogeneously located throughout the periphery of the cells (Aguilar et al., 2011; Carranza et al., 2021). Instead, the observed lateral localisation of active conjugation pores may reflect the facilitated access to F plasmid molecules, which are also positioned at quarter positions and excluded from the cell poles (Gordon et al., 2004; Niki and Hiraga, 1997). By contrast, the ssDNA mainly enters the polar region of the recipient cells. This could suggest that the pole of the recipients’ surface is the preferred location for the donor’s F-pilus attachment or the stabilisation of the mating pair. The latter possibility is reinforced by the fact that mating pair stabilisation during F conjugation involves interaction between the plasmid protein TraN exposed at the surface of the donor cells and the host outer membrane protein OmpA of the recipient cells (Klimke and Frost, 1998; Low et al., 2022). OmpA was shown to be enriched and less mobile in the polar regions of *E. coli* cells (Verhoeven et al., 2013), possibly favouring the stabilisation of the mating pair and the conjugation pore at this location.

The unexpected finding that the ssDNA is present in the donor during conjugation also provides insights into the activity of TraI and its coordination with the transfer of the T-strand through the T4SS or the RCR of the non-transferred strand. Before DNA transfer initiation, the relaxosome bound to the plasmid’s *oriT* is docked to the T4SS by the TraD (VirD4) coupling protein, thus forming the pre-initiation complex (Figure 5A(i)). Contact with the recipient cell is proposed to induce a signal that activates the pre-initiation complex. We uncover the existence of a brief stage where part of the T-strand has already been transferred into the recipient cell while no ssDNA is present within the donor (Figure 5A(ii). At this stage, the absence of ssDNA in the donor implicates that all the ssDNA generated by TraI has been removed, both by transfer of the T-strand through the T4SS and by complementation of the non-transferred ssDNA strand by the RCR. After this transient stage, the ssDNA also accumulates in the donor, suggesting that the ssDNA is generated by TraI helicase activity in the donor faster than it is removed by transfer and RCR synthesis (Figure 5A(iii).

**Figure 5.**
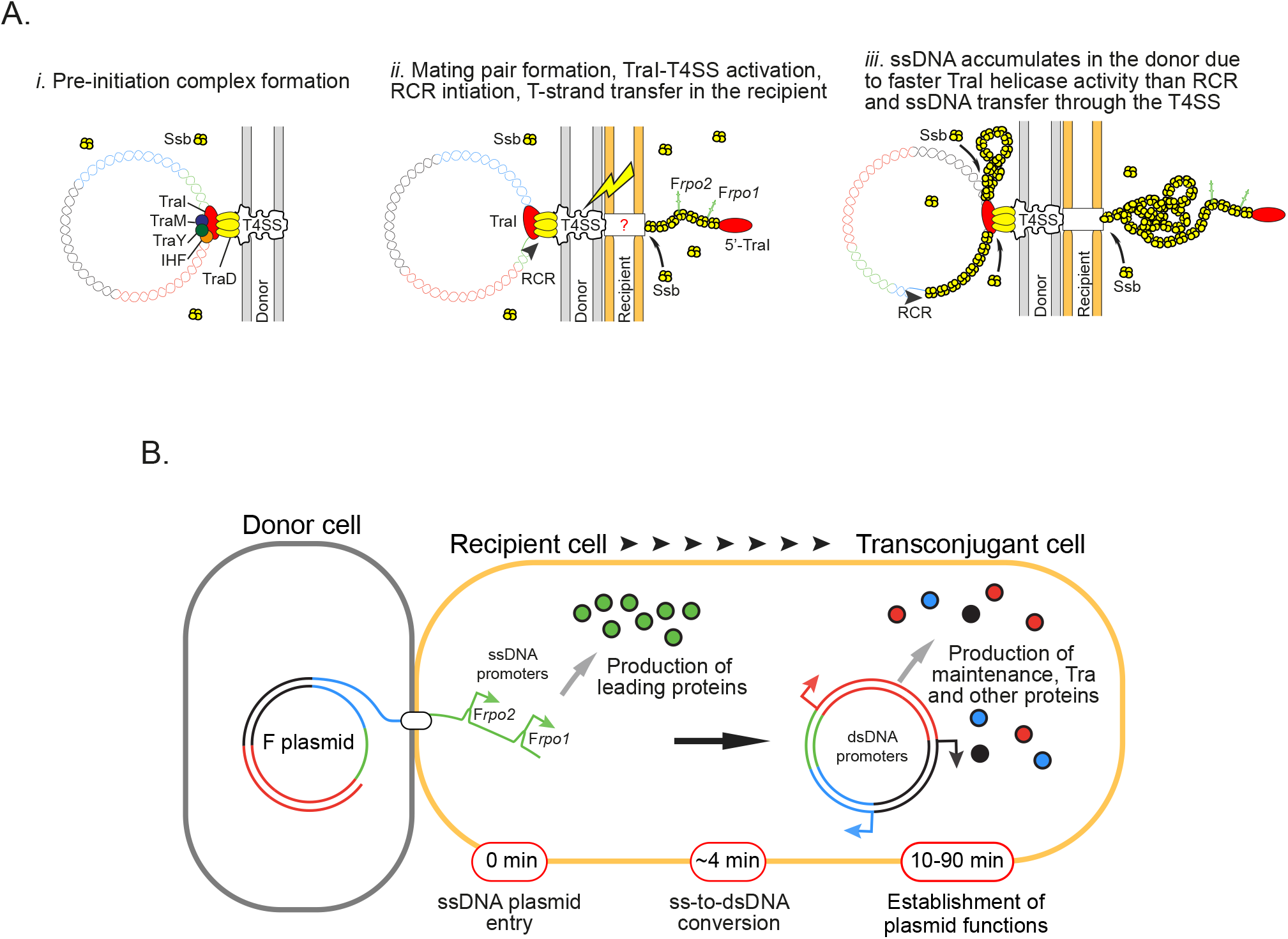
Model for conjugation initiation and intracellular dynamics. **(A)** (*i*) Before the initiation of conjugation, the pre-initiation complex bound to the plasmid’s origin of transfer is docked to the Type IV secretion system (T4SS). (*ii*) The establishment of the mating pair transduces a signal that activates the pre-initiation complex. Unwinding of the dsDNA plasmid by the helicase activity of TraI produces the first segment of the T-strand, which is immediately transferred into the recipient cell where it recruits Ssb molecules, while the non-transferred strand is being complemented by rolling-circle replication (RCR) in the donor cell. (*iii*) The helicase activity of TraI generates ssDNA at higher rate than the T-strand is transferred through the T4SS or the non-transferred strand is complemented by RCR, thus resulting in the accumulation of ssDNA plasmid coated by Ssb molecules in the donor cell. **(B)** Upon entry of the ssDNA plasmid in the recipient cell, F*rpo1* and F*rpo2* leading sequences form stem-loop structures that serve as promoters initiating the transcription of the downstream leading genes, rapidly resulting in the production of leading proteins. The subsequent ss-to-dsDNA conversion inactivates F*rpo1* and F*rpo2* and licenses the expression of other plasmid genes under the control of conventional dsDNA promoters. The production of maintenance, transfer and other plasmid-encoded proteins eventually results in the development of new functions by the transconjugant cell.

Assuming the 2.9 ± 1.1 min lifespan of the Ssb-Ypet foci in transconjugants reflects the time required to complete the internalisation of the 108 000 nt ssDNA F plasmid, we calculated a 620 ± 164 nt.s^-1^ transfer rate. This is in reasonable agreement with the historical 770 nt.s^-1^ rate estimated from the 100 minutes required to transfer the whole 4.6 Mb *E. coli* chromosome (Jacob and Wollman, 1958). Besides, the rate of DNA synthesis by the DNA polymerase III holoenzyme during RCR was estimated at 650-750 nuc.s^-1^ (Stephens and McMacken, 1997). By comparison, the rate of TraI helicase activity was measured at 1120 ± 160 bp.s^-1^ (Sikora et al., 2006). These estimates support the view that ssDNA accumulation in the donor accounts for the faster rate of TraI helicase activity than the rate of T-strand plasmid transfer or RCR. Therefore, it is possible that, contrasting with the previously suggested but never demonstrated proposal, the helicase activity of the relaxase is not strictly coupled with the activity of DNA translocation through the T4SS.

Live-cell microscopy uncovers the global chronology conjugation steps, as summarised in Figure 5B. The plasmid processing in the transconjugant cell is a relatively rapid process, as the entry of the ssDNA plasmid and its conversion into dsDNA is completed in about 4 minutes on average. Most importantly, the ss-to-dsDNA conversion event is the pivotal event that determines the program of plasmid gene expression. Leading genes are the first to enter the recipient cell and also the first to be expressed from the F plasmid in ssDNA form. Consistently with previous proposals (Bates et al., 1999; Masai and Arai, 1997; Nasim et al., 2004), we show that the early expression of leading genes depends on sequences that act as single-stranded promoters when the plasmid is still in ssDNA form. As previously described for F*rpo1*, we propose that the highly homologous F*rpo2* sequences identified here folds into a stable stem-loop structures that reconstruct -35 and -10 consensus boxes, resulting in transcription initiation.

Leading gene expression is also transient as the ss-to-dsDNA conversion turns off leading protein production by inactivating F*rpo1* and F*rpo2* promoters while licencing the expression of maintenance, transfer and other plasmid genes under the control of conventional dsDNA promoters, often subject to their own regulation specificities. Maintenance and transfer protein levels within transconjugants reach a steady-state equivalent to that of vegetatively growing F-containing cells in about 30 to 90 minutes, depending on the protein. Interestingly, our previous work showed that tetracycline resistance factors encoded by the Tn*10* transposon inserted in the intergenic region *ybdB-ybfA* of the F plasmid are also produced immediately after the ss-to-dsDNA conversion and reach the resistant cell’s level within approximately 90 minutes (Nolivos et al., 2019). These findings consistently indicate that this time scale corresponds to the period needed for the transconjugant cells to gain plasmid-encoded functions, including plasmid maintenance, conjugation ability, immunity against self-transfer and additional resistance potentially carried by the plasmid.

The regulation of plasmid gene expression by plasmid processing is an elegant way to ensure the sequential and timely production of plasmid proteins in the transconjugant cell, and particularly to restrict the production of leading factors to a narrow time window following the entry of the ssDNA plasmid. However, d*e novo* protein synthesis might not be the only way to provide the transconjugant cell with plasmid-encoded proteins. Recent work by Al Mamun et *al*. reports that the transfer of the F-like plasmid pED208 (IncFV) is concomitant with the translocation of several plasmid-encoded proteins, including TraI, ParA, ParB1, Ssb homologue Ssb^ED208^, ParB2, PsiB, and PsiA (Al Mamun et al., 2021). Protein translocation was detected at low frequency (10^−5^ recombinants per donor cell between one and five hours of mating) using a highly sensitive Cre recombinase assay. Protein translocation might also occur during the transfer of the native F plasmid but could not solely explain our observations. Indeed, our microscopy analysis shows that YgeA, PsiB, YfjB, YfjA and Ssb^F^ leading fusions are below the microscopy detection threshold in donor cells but are quantified at significant intracellular levels in all transconjugant cells. This implies that the amounts of leading proteins observed in the transconjugant cells cannot just originate from donor cells, but result from *de novo* protein synthesis, which we show depends on F*rpo1* and F*rpo2* sequences.

Both the early production and the direct translocation of leading proteins suggest a critical role of the leading region in conjugation. Several elements support this view. The leading region is conserved in a variety of conjugative plasmids (Cox and Schildbach, 2017; Golub and Low, 1985, 1986a; Golub et al., 1988; Loh et al., 1989, 1990). In addition, the leading regions of plasmids belonging to a wide range of incompatibility groups (IncF, IncN, IncP9 and IncW) classified as MOBF plasmids using the relaxase as a phylogenetic marker were reported to be the preferential target for CRISPR-Cas systems directed against conjugation (Fernandez-Lopez et al., 2016; Garcillán-Barcia et al., 2009; Westra et al., 2013). Recently, the leading region was shown to be an important evolutionary target for the dissemination of the pESLB (IncI) plasmid (Benz and Hall, 2022). Concerning the F plasmid, we can stress that F*rpo1* and F*rpo2* share 92 % similarity at the nucleotide level and are located only about 5 kb apart. This implies that when in dsDNA form during vegetative plasmid replication, F*rpo1* and F*rpo2* sequences would be a potential substrate for homologous recombination, resulting in the deletion of the intervening segment. However, the intervening segment carries the *flmAB* genes, functional homologues to the *hok/sok* toxin-antitoxin system from the R1 plasmid (Loh et al., 1988), which are likely to safeguard the stability of the leading region.

Despite this body of evidence, it is currently challenging to rationalise the importance of the leading region since the molecular functions of most leading proteins are still unknown. Our data indicate that genes downstream of F*rpo1* (*ygeA* et *ygeB*) have a critical function in conjugation. By contrast, genes located downstream F*rpo2* (*ssb*^*F*^, *yfjA, yfjB, psiB, psiA* and *flmC*) appear to be dispensable since deletions of F*rpo2, ssb*^*F*^ or *psiB* (Loh et al., 1989) have no significant impact on the overall conjugation efficiency addressed by plating assays. However, conjugation efficiency assays are generally performed between identical or closely related bacterial strains in optimal medium and temperature conditions. This likely undermines the role of genes that are not strictly essential but might facilitate or optimise conjugation. Hence, it is possible that the importance of the leading factors would be best revealed in less favourable conditions, between phylogenetically distant bacteria, or on the evolutionary scale. Meanwhile, real time microscopy might help uncover the potentially subtle influence of these genes on the sequence of conjugation in live cells.

## Acknowledgements

The authors thank the National BioResource Project and Coli Genetic Stock Center for providing strains, A. Ducret for valuable help with MicrobeJ and N. Fraikin for helpful discussion.

## Funding

This research was funded by the Foundation for Medical Research, grant number FRM-EQU202103012587 to C.L. and A.C.; the French National Research Agency, grant number ANR-18-CE35-0008 to C.L., Y.Y., and K. G.; and the University of Lyon through funding to C.V. C.L. also acknowledges the Schlumberger Foundation for Education and Research (FSER 2019).

## Author contributions

C.L. and S.B. conceived, designed and supervised the execution of the study; A.C., C.V., K.G., A.R., S.N. and S.B. performed the experiments and analysed the data. C.L. and S.B. wrote the paper, and C. L. prepared the figures. C.L. and Y.Y. provided funding.

## Competing Financial Interests

The authors declare no competing financial interests.

## Data and materials availability

All data to understand and assess the conclusions of this research are available in the main text and Supplementary Materials.

## Materials and Methods

### Bacterial strains, plasmids and growth

Bacterial strains are listed in Table S1, plasmids in Table S2, and oligonucleotides in Tables S3. Fusion of genes with fluorescent tags and gene deletion on the F plasmid used λRed recombination (Datsenko and Wanner, 2000; Yu et al., 2000). Modified F plasmids were transferred to the background strain K12 MG1655 by conjugation. Where multiple genetic modifications on the F plasmid were required, the *kan* and *cat* genes were removed using site-specific recombination induced by expression of the Flp recombinase from plasmid pCP20 (Datsenko and Wanner, 2000). Plasmid cloning were done by Gibson Assembly and verified by Sanger sequencing (Eurofins Genomics biotech). Strains and plasmids were verified by Sanger sequencing (Eurofins Genomics). Cells were grown at 37°C in M9 medium supplemented with glucose (0.2 %) and casamino acid (0.4 %) (M9-CASA) before imaging, and in Luria-Bertani (LB) broth for conjugation efficiency assays. When appropriate, supplements were used in the following concentrations; Ampicillin (Ap) 100 μg/ml, Chloramphenicol (Cm) 20 μg/ml, Kanamycin (Kn) 50 μg/ml, Streptomycin (St) 20 μg/ml, and Tetracycline (Tc) 10 μg/ml.

### Conjugation assays

Overnight cultures in LB of recipient and donor cells were diluted to an A_600_ of 0.05 and grown until an A_600_ comprised between 0.7 and 0.9 was reached. 25 μl of donor and 75 μl of recipient cultures were mixed into an Eppendorf tube and incubated for 90 minutes at 37°C. 1 ml of LB was added gently and the tubes were incubated again for 90 min at 37°C. Conjugation mix were vortexed, serial diluted, and plated on LB agar X-gal 40 μg/ml IPTG 20 μM supplemented the appropriate antibiotic to select for recipient or donor populations. Recipient (R) colonies were then streaked on plated on LB agar containing tetracycline 10 μg/ml to select for transconjugants (T) and the frequency of transconjugant calculated from the (T/R+T) presented in Figure 4C.

### Live-cell microscopy experiments

Overnight cultures in M9-CASA were diluted to an A_600_ of 0.05 and grown until A_600_ = 0.8 was reached. Conjugation samples were obtained by mixing 25 μl of donor and 75 μl of recipient into an Eppendorf tube. For time-lapse experiments, 50 μl of the pure culture or conjugation mix was loaded into a B04A microfluidic chamber (ONIX, CellASIC®) (Cayron and Lesterlin, 2019). Nutrient supply was maintained at 1 psi and the temperature maintained at 37°C throughout the imaging process. Cells were imaged every 1 or 5 min for 90 to 120 minutes. For snapshot imaging, 10 μl samples of clonal culture or conjugation mix were spotted onto an M9-CASA 1% agarose pad on a slide (Lesterlin and Duabrry, 2016) and imaged directly.

#### Image acquisition

Conventional wide-field fluorescence microscopy imaging was carried out on an Eclipse Ti2-E microscope (Nikon), equipped with x100/1.45 oil Plan Apo Lambda phase objective, ORCA-Fusion digital CMOS camera (Hamamatsu), and using NIS software for image acquisition. Acquisitions were performed using 50% power of a Fluo LED Spectra X light source at 488 nm and 560 nm excitation wavelengths. Exposure settings were 100 ms for Ypet, sfGFP and mCherry and 50 ms for phase contrast.

#### Image analysis

Quantitative image analysis was done using Fiji software with MicrobeJ plugin (Ducret et al., 2016). For snapshot analysis, cells’ outline detection was performed automatically using MicrobeJ and verified using the Manual-editing interface. For time-lapse experiments, detection of cells was done semi-automatedly using the Manual-editing interface, which allows to select the cells to be monitored and automatically detect the cell outlines. Within conjugation populations, donor (no mCh-ParB signal), recipient (diffuse mCh-ParB signal), or transconjugant (mCh-ParB foci) category were assigned using the ‘Type’ option of MicrobeJ. Recipient cells were detected on the basis of the presence of red fluorescence above the cell’s autofluorescence background level detected in the donors. Among these recipient cells, transconjugants were identified by running MicrobeJ automated detection of the ParB fluorescence foci (Maxima detection). This approach was used independently of the presence or the absence of the Ssb-Ypet, or sfGFP fusions within donor and recipient cells. Within the different cell types, mean intensity fluorescence (a.u.), skewness, Signal/Noise Ratio (SNR), or cell length (μm) parameters were automatically extracted and plotted using MicrobeJ. SNR corresponds to the ratio (mean intracellular signal / mean noise signal), where the mean intracellular signal is the fluorescence signal per cell area and the noise is the signal measured outside the cells (due to the fluorescence emitted by the surrounding medium). By contrast with the total amount of fluorescence per cell, which is depending on the cell size/age and accounts for the background, SNR quantitative estimate is more appropriate for unbiased quantification of intracellular fluorescence over time. Ssb-Ypet, Ssb^F^-mCh and mCh-ParB foci were detected using MicrobeJ Maxima detection function, and foci localisation and fluorescence intensity were extracted and plotted automatically. Plots presenting time-lapse data were either aligned to the first frame where the transconjugant cell exhibits a conjugative Ssb-Ypet focus (ssDNA acquisition) or a mCh-ParB focus (ss-to-dsDNA conversion) as indicated in the corresponding figure legend.

## Statistical analysis

*P-*value significance were analysed running specific statistical tests on the GraphPad Prism software. Single-cell data from quantitative microscopy analysis were extracted from the MicrobeJ interface and transferred to GraphPad. P-value significance of single-cell quantitative data was performed using unpaired non-parametric Mann-Whitney statistical test, which allows to compare differences between independent data groups without normal distribution assumption. *P-*value significance for the frequency of transconjugants obtained by plating assays were evaluated using One-way analysis of variance (ANOVA) with Dunnetts multiple comparisons test, which allows to determine the statistical significant of differences observed between the means of three or more independent [experimental groups](https://www.statisticshowto.com/experimental-group/) against a control group mean (corresponding to the F*wt*). When required, *P-* value and significance are indicated on the figure panels and within the corresponding legend.

## SUPPLEMENTARY MATERIALS

Figs. S1 to S7

Tables S1 to S3

Captions for Movies S1 to S3

Movies S1 to S3

## Notes

### Competing Interest Statement

The authors have declared no competing interest.

