## Supplementary material for "Real time visualisation of conjugation reveals the molecular strategy evolved by the conjugative F plasmid to ensure the sequential production of plasmid factors during establishment in the new host cell": Supplmentary Material

**This file includes:**

Figures S1 to S7
Tables S1 to S3
Captions for Movies S1 to S3

**Other Supplementary Materials for this manuscript include the following:**

Movies S1 to S3

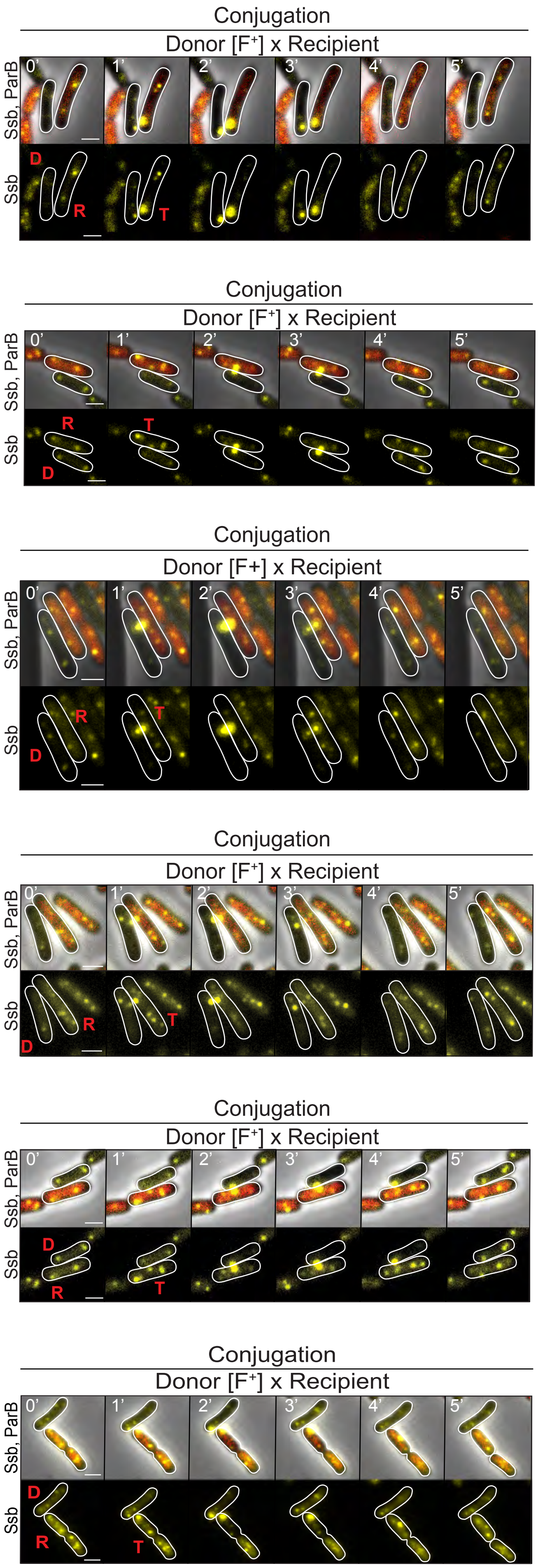
Conjugation
Donor [F⁺] x Recipient
Ssb, ParB
Ssb
0'
1'
2'
3'
4'
5'
D
R
T
Conjugation
Donor [F⁺] x Recipient
Ssb, ParB
Ssb
0'
1'
2'
3'
4'
5'
R
T
D
Conjugation
Donor [F+] x Recipient
Ssb, ParB
Ssb
0'
1'
2'
3'
4'
5'
R
T
D
Conjugation
Donor [F⁺] x Recipient
Ssb, ParB
Ssb
0'
1'
2'
3'
4'
5'
R
T
D
Conjugation
Donor [F⁺] x Recipient
Ssb, ParB
Ssb
0'
1'
2'
3'
4'
5'
D
R
T
Conjugation
Donor [F⁺] x Recipient
Ssb, ParB
Ssb
0'
1'
2'
3'
4'
5'
D
R
T

Figure S1

**Figure S1. ssDNA transfer events reported by the formation of Ssb-Ypet conjugative foci.** Time-lapse microscopy images of conjugation performed in microfluidic chamber showing a plasmid transfer event between a donor (D) and a recipient cell (R) that is converted into a transconjugant (T) (similar to Figure 1B). The transfer of the ssDNA plasmid is reported by the formation of paired bright membrane-associated Ssb-Ypet conjugative foci in both the donor and the transconjugant cell. The recipient cells also produce mCh-ParB. Scale bars 1 µm. Donor (LY1007), recipient (LY358), transconjugant (LY358 after F*wt* acquisition from LY1007).

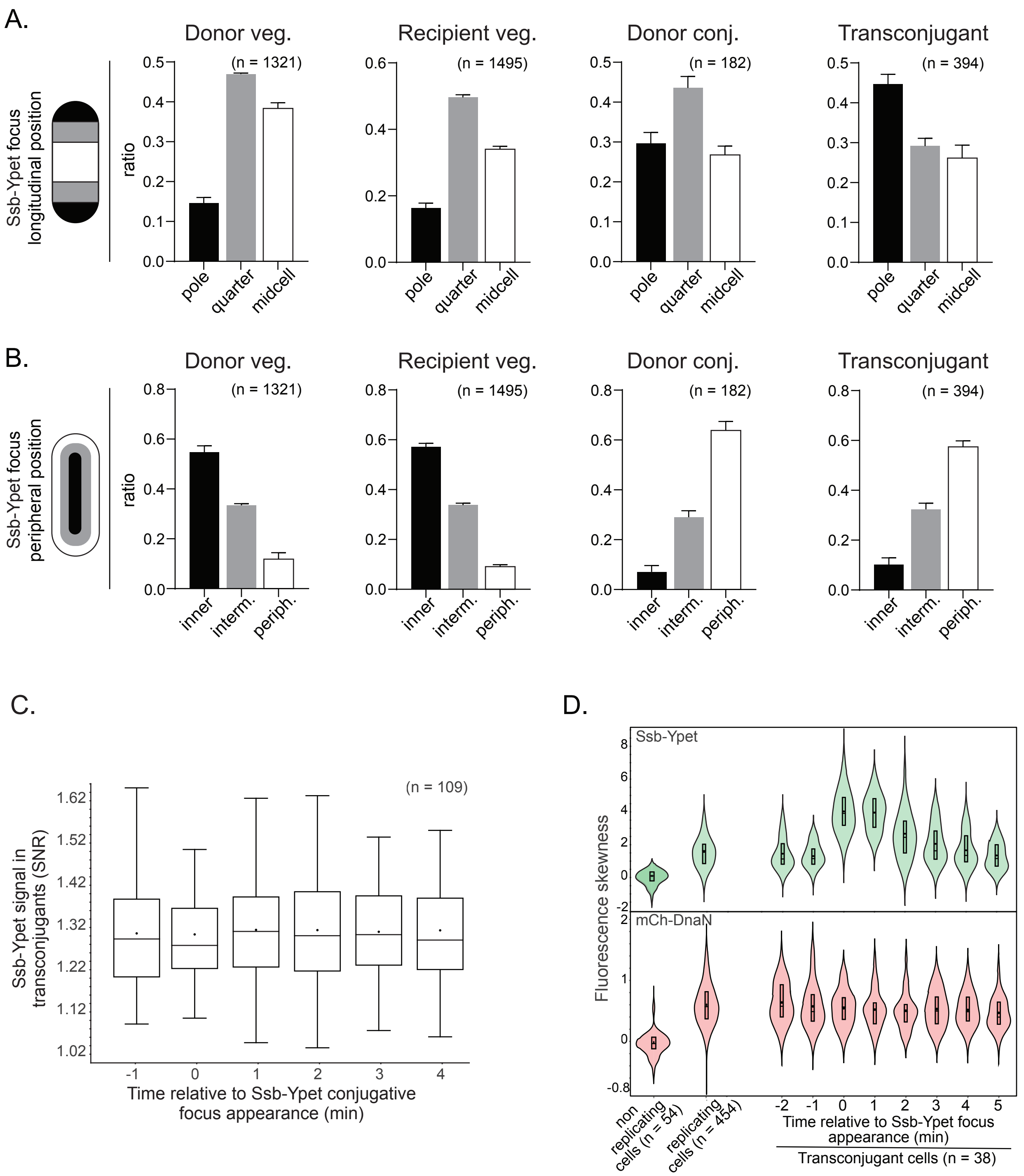
A.
Ssb-Ypet focus longitudinal position
Donor veg.
(n = 1321)
Recipient veg.
(n = 1495)
Donor conj.
(n = 182)
Transconjugant
(n = 394)
ratio
pole
quarter
midcell
B.
Ssb-Ypet focus peripheral position
inner
interm.
periph.
C.
(n = 109)
Ssb-Ypet signal in transconjugants (SNR)
Time relative to Ssb-Ypet conjugative focus appearance (min)
D.
Ssb-Ypet
mCh-DnaN
Fluorescence skewness
non replicating cells (n = 54)
replicating cells (n = 454)
Time relative to Ssb-Ypet focus appearance (min)
Transconjugant cells (n = 38)

Figure S2

**Figure S2. Analysis of Ssb-Ypet foci localisation in donor and recipient cells during vegetative growth or during conjugation.**

(**A**) Histograms showing the localisation distribution of Ssb-Ypet foci along the cell length sorted in polar, quarter and midcell categories as depicted on the cell diagram (left). (**B**) Histograms showing the localisation distribution of Ssb-Ypet foci within the cell compartment sorted in inner, intermediate and peripheral (i.e., membrane proximal) categories as depicted on the cell diagram (left). For (**A**) and (**B**), the analysis was performed on the same data set as for Figure 1C-D and 1H. The number (n) of cells analysed is indicated. F*wt* Donor (LY1007), Recipient (LY358) (**C**) Boxplot showing stable Ssb-Ypet intracellular fluorescence (SNR) in transconjugant cells with respect to the ssDNA transfer initiated at t = 0 min. The median, quartile 1 and quartile 3 are indicated by horizontal lines and the mean by a black dot. The analysis was performed on the same data set than for Figure 1I. The number (n) of transconjugant cells analysed is indicated. Transconjugants (LY358 after F*wt* acquisition from LY1007). (D) Violin plot showing the skewness of mCh-DnaN and Ssb-Ypet fluorescence in non-replicating and replicating recipient cells in vegetative growth, and in transconjugants with respect to the ssDNA transfer initiated at t = 0 min, as reflected by Ssb-Ypet skewness increase. The number (n) of cells analysed is indicated. Recipient (LY355), transconjugants (LY355 after F*wt* acquisition from LY1007).

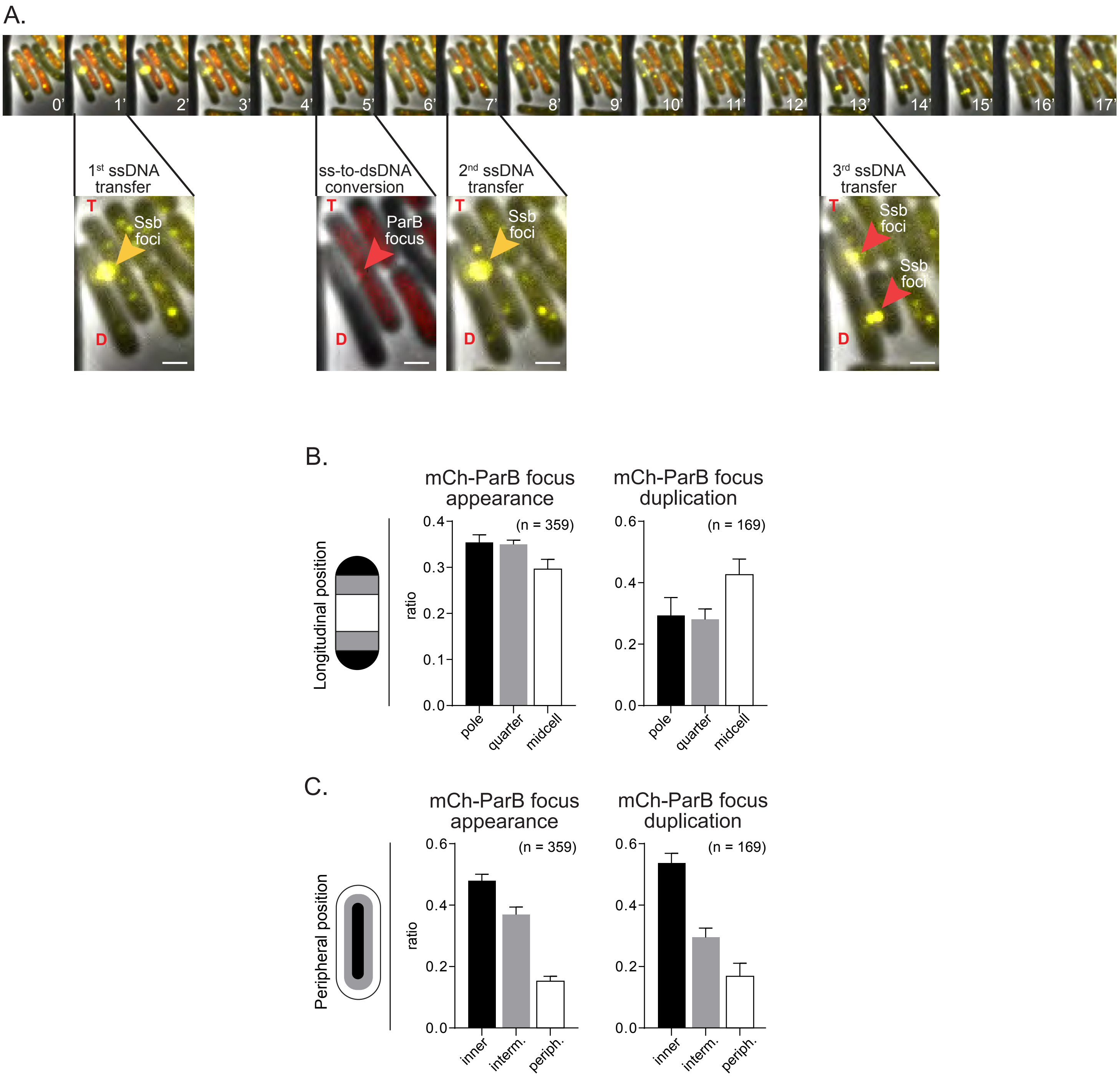
A.
0'
1'
2'
3'
4'
5'
6'
7'
8'
9'
10'
11'
12'
13'
14'
15'
16'
17'
1st ssDNA transfer
ss-to-dsDNA conversion
2nd ssDNA transfer
3rd ssDNA transfer
T
D
Ssb foci
ParB focus
B.
Longitudinal position
mCh-ParB focus appearance
(n = 359)
mCh-ParB focus duplication
(n = 169)
ratio
pole
quarter
midcell
C.
Peripheral position
mCh-ParB focus appearance
(n = 359)
mCh-ParB focus duplication
(n = 169)
ratio
inner
interm.
periph.

Figure S3

**Figure S3. Multiple ssDNA transfer and mCh-ParB localisation analysis.**

(**A**) Time lapse microscopy images showing a transconjugant cell that acquires multiple ssDNA plasmids reflected by three consecutive appearances of Ssb-Ypet conjugative foci in both the donor (D) and the transconjugant (T) cells. The transconjugant acquires a first ssDNA plasmid (t1 min) followed by the appearance of the mCh-ParB focus that reflects the successful ss-to-dsDNA conversion (t5 min), then a second (t7 min) and a third (t=13 min) ssDNA plasmid. All three ssDNA come from the same donor cell and appear to occur at the same position along the cell membrane. Scale bars 1 µm. (**B**) and (**C**) Histograms showing the localisation distribution of mCh-ParB foci at the time of appearance and just before its duplication into two foci in the transconjugant cells. The analysis was performed on the same data set than for Figure 2H. (**B**) Localisation have been sorted according their polar, quarter and midcell position along the cell length, as depicted on the cell diagram (left). (**C**) Localisation have been sorted according to their inner, intermediate and peripheral (i.e., membrane proximal) positions, as depicted on the cell diagram (left). Donor (LY1007), Recipient (LY358).

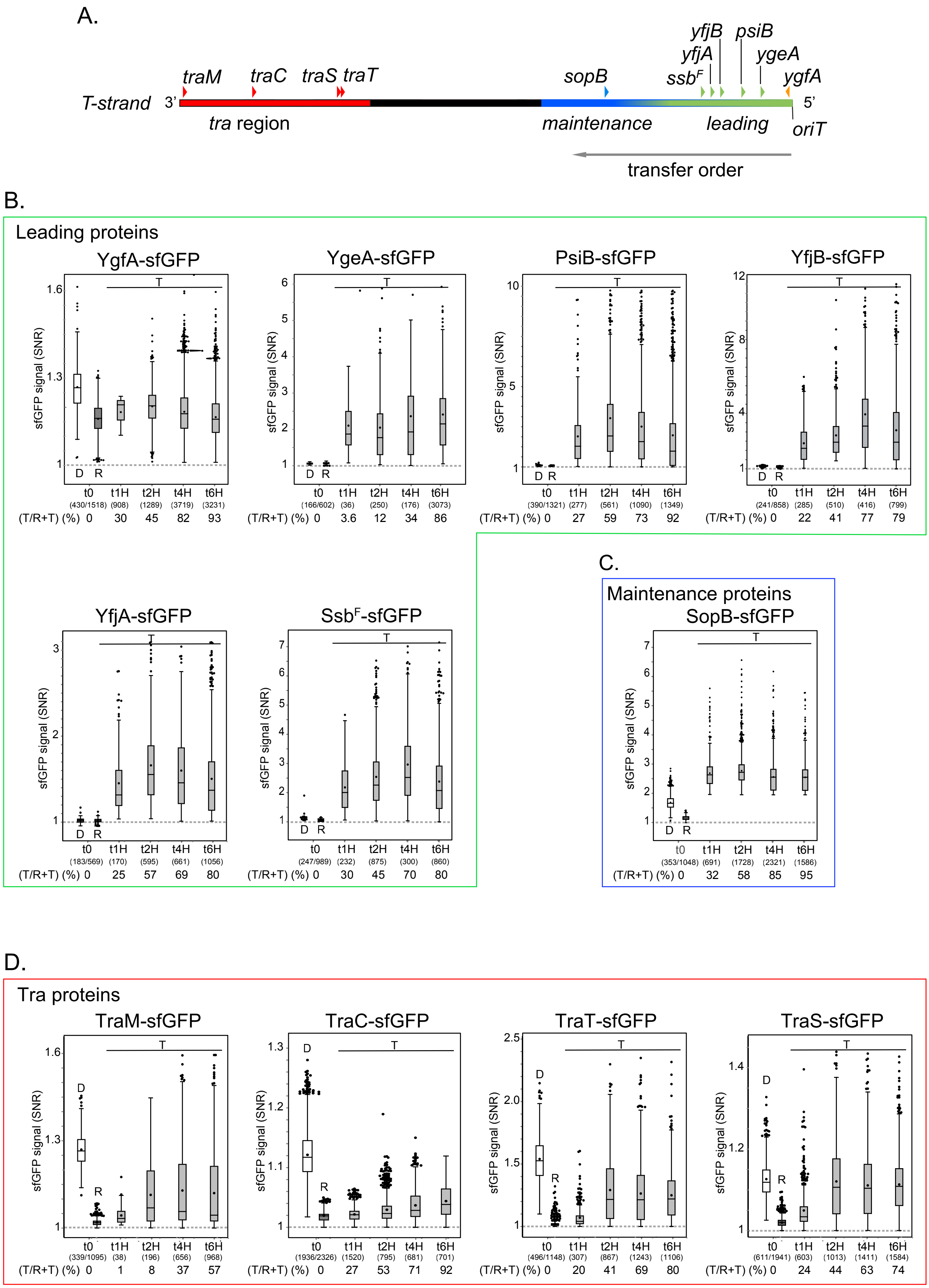
A.
traM
traC
traS traT
sopB
yfjB
psiB
yfjA
ygeA
ssbF
ygfA
T-strand
3'
5'
tra region
maintenance
leading
oriT
transfer order
B.
Leading proteins
YgfA-sfGFP
YgeA-sfGFP
PsiB-sfGFP
YfjB-sfGFP
sfGFP signal (SNR)
D
R
T
t0
t1H
t2H
t4H
t6H
(430/1518) (908) (1289) (3719) (3231)
(T/R+T) (%) 0 30 45 82 93
(166/602) (36) (250) (176) (3073)
(T/R+T) (%) 0 3.6 12 34 86
(390/1321) (277) (561) (1090) (1349)
(T/R+T) (%) 0 27 59 73 92
(241/858) (285) (510) (416) (799)
(T/R+T) (%) 0 22 41 77 79
YfjA-sfGFP
SsbF-sfGFP
(183/569) (170) (595) (661) (1056)
(T/R+T) (%) 0 25 57 69 80
(247/989) (232) (875) (300) (860)
(T/R+T) (%) 0 30 45 70 80
C.
Maintenance proteins
SopB-sfGFP
(353/1048) (691) (1728) (2321) (1586)
(T/R+T) (%) 0 32 58 85 95
D.
Tra proteins
TraM-sfGFP
TraC-sfGFP
TraT-sfGFP
TraS-sfGFP
(339/1095) (38) (196) (656) (968)
(T/R+T) (%) 0 1 8 37 57
(1936/2326) (1520) (795) (681) (701)
(T/R+T) (%) 0 27 53 71 92
(496/1148) (307) (867) (1243) (1106)
(T/R+T) (%) 0 20 41 69 80
(611/1941) (603) (1013) (1411) (1584)
(T/R+T) (%) 0 24 44 63 74

Figure S4

**Figure S4. Time-course analysis of sfGFP fusion production in transconjugant cells.**

(**A**) Genetic map of the F plasmid showing the position, orientation and order of transfer from the *oriT*, of the genes for which C-terminal translational sfGFP fusions have been studied. (**B**), (**C**) and (**D**), box plots showing the quantification of sfGFP fusions intracellular signal (SNR) during conjugation time-course experiments. For each plot, data are presented for donor (D) and recipient (R) cells at t = 0 min, and transconjugant (T) cells at 1, 2, 4 and 6 hours after mixing donors and recipient cells. The median, quartile 1 and quartile 3 are indicated by horizontal lines and the mean by a black dot. Black dots above and below the max and min values correspond to outlier cells. The number of cells analysed (n) from three biological replicates is indicated between brackets, as well as the frequency of transconjugants (T/R+T) directly measured at the single-cell level from the proportion of recipient cells exhibiting diffuse mCh-ParB fluorescence (R) or transconjugant cells harbouring mCh-ParB foci (T). Results are presented for the leading (**B**), maintenance (**C**) and *tra* genes (**D**). Donor (see Table S1), Recipient (LY358).

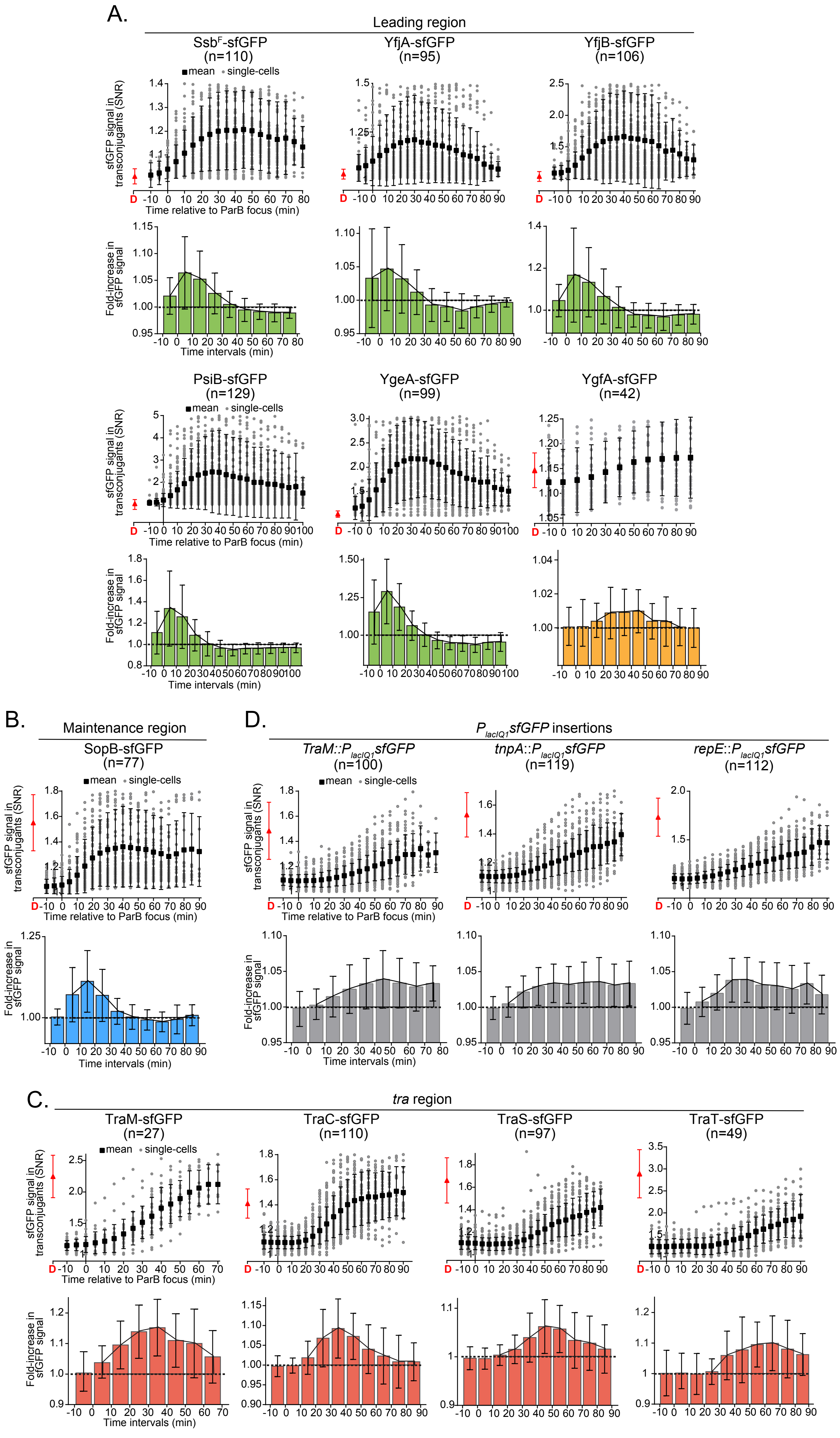
A.
Leading region
SsbF-sfGFP (n=110)
YfjA-sfGFP (n=95)
YfjB-sfGFP (n=106)
PsiB-sfGFP (n=129)
YgeA-sfGFP (n=99)
YgfA-sfGFP (n=42)
mean
single-cells
sfGFP signal in transconjugants (SNR)
Time relative to ParB focus (min)
Fold-increase in sfGFP signal
Time intervals (min)
D
B.
Maintenance region
SopB-sfGFP (n=77)
D.
PlacIQ1sfGFP insertions
TraM::PlacIQ1sfGFP (n=100)
tnpA::PlacIQ1sfGFP (n=119)
repE::PlacIQ1sfGFP (n=112)
C.
tra region
TraM-sfGFP (n=27)
TraC-sfGFP (n=110)
TraS-sfGFP (n=97)
TraT-sfGFP (n=49)

Figure S5

**Figure S5. Time lapse analysis of sfGFP fusion production in transconjugant cells.**

(**A-D**) The production level of sfGFP with respect to the appearance of the mCh-ParB focus in individual transconjugant cells was analysed from time lapse microscopy imaging of conjugation performed in microfluidic chambers. Results are presented for the leading (**A**), maintenance (**B**) *tra* genes (**C**), and $P_{lacIQ1}$sfGFP reporter insertions (**D**). For each fusion, the top panel shows the quantification of sfGFP intracellular signal (SNR) in transconjugant cells over time with respect to mCh-ParB focus appearance (t = 0 min). Each data point representing a single cell (grey dots), the mean (black square) and the SD from the indicated (n) number of transconjugant cells analysed are shown. The mean SNR and SD observed in donor cells during vegetative growth is also shown (red) for comparison. The bottom panel presents the fold-increase in sfGFP SNR signal by 10 minutes interval calculated from the same data set. Fold-increase >1 reveals the production of sfGFP by the transconjugants in the corresponding time interval. The black polygons show the data that have been used to generate Figure 3B. Donor (see Table S1), Recipient (LY318).

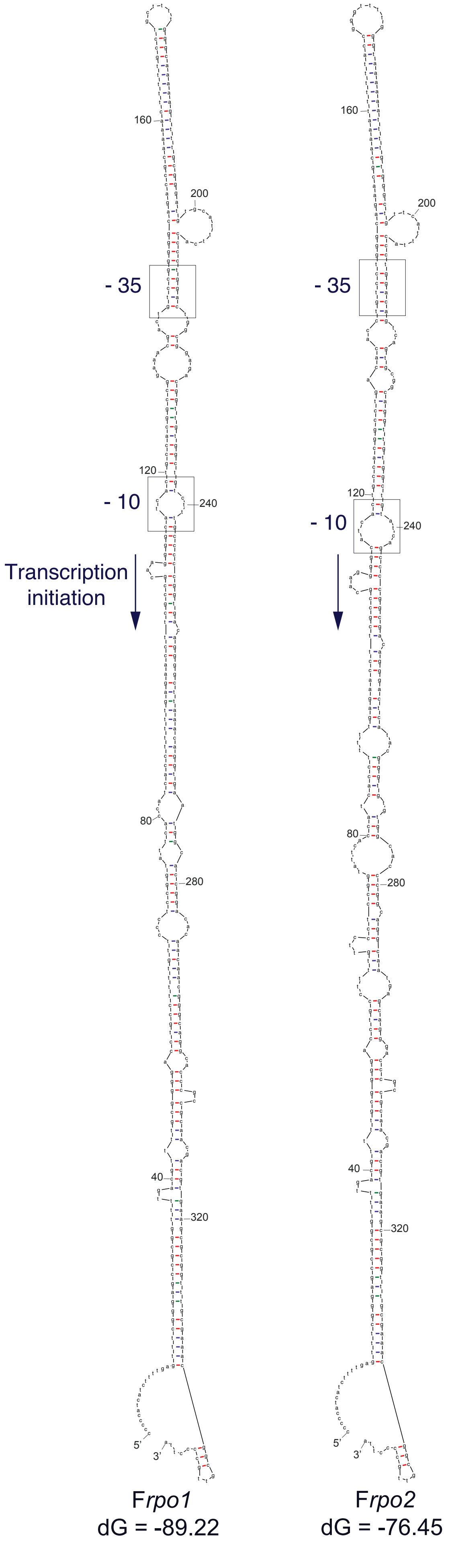
160
200
- 35
120
- 10
240
Transcription initiation
80
280
40
320
5'
3'
Frpo1
dG = -89.22
160
200
- 35
120
- 10
240
80
280
40
320
5'
3'
Frpo2
dG = -76.45

Figure S6

**Figure S6. F*rpo1* and F*rpo2* stem-loop structure.**

Secondary structure folding prediction for F*rpo1* and F*rpo2* sequences in ssDNA forms generated by mFOLD. The reconstructed -35 and -10 boxes are shown (black boxes) with the predicted orientation of transcription initiation. The Gibbs free energy change (dG) indicative of the structure's thermodynamic stability is indicated.

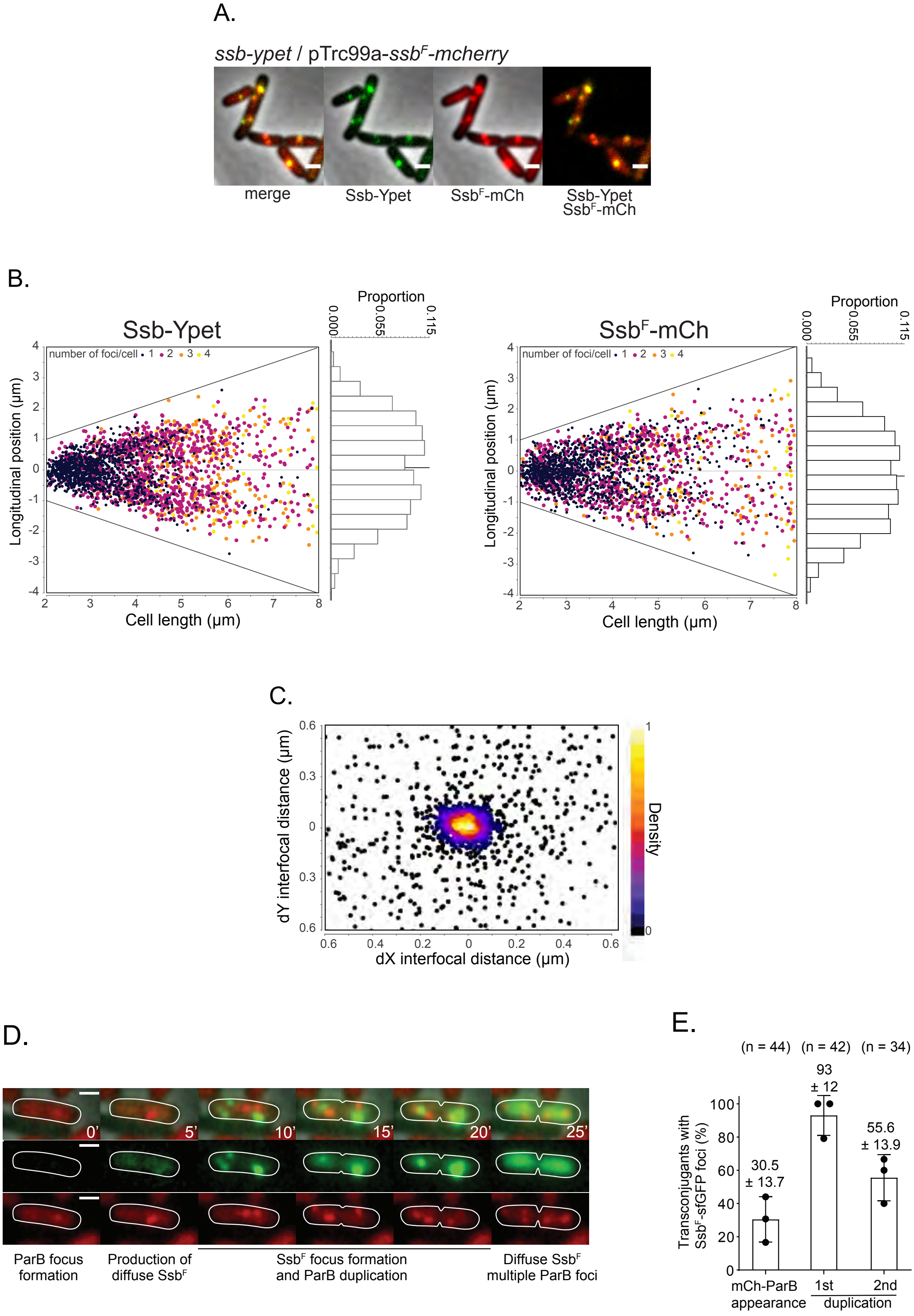
A.
ssb-ypet / pTrc99a-ssbF-mcherry
merge
Ssb-Ypet
SsbF-mCh
Ssb-Ypet SsbF-mCh
B.
Ssb-Ypet
SsbF-mCh
Proportion
0.000
0.055
0.115
number of foci/cell • 1 • 2 • 3 • 4
Longitudinal position (µm)
Cell length (µm)
C.
dY interfocal distance (µm)
dX interfocal distance (µm)
Density
D.
0'
5'
10'
15'
20'
25'
ParB focus formation
Production of diffuse SsbF
SsbF focus formation and ParB duplication
Diffuse SsbF multiple ParB foci
E.
(n = 44)
(n = 42)
(n = 34)
93 ± 12
30.5 ± 13.7
55.6 ± 13.9
Transconjugants with SsbF-sfGFP foci (%)
mCh-ParB appearance
1st
2nd
duplication

Figure S7

**Figure S7. Analysis of Ssb-Ypet and Ssb$^F$-mCh localisation**

**(A)** Microscopy images showing the colocalisation between the chromosomally encoded Ssb-Ypet and Ssb$^F$-mCh ectopically produced from an expression plasmid during vegetative growth. Scale bars 1µm. Strain LY1382. (**B**) Position analysis of Ssb-Ypet (left) and Ssb$^F$-mCh (right) foci during vegetative growth presented as localisation dot plots and histogram of distribution along the cell length. (**C**) 2D colocalisation density plot presenting the interfocal distance between Ssb-Ypet and Ssb$^F$-mCh foci during vegetative growth. The positions of Ssb-Ypet foci are normalised at coordinate (dX = 0, dY = 0) to serve as a localisation reference. The density scale is shown. (**D**) Time lapse microscopy images of conjugation performed in microfluidic chamber showing the formation of mCh-ParB focus (t = 0 min) followed by the production of diffuse Ssb$^F$-sfGFP (t = 5 min), and formation Ssb$^F$-sfGFP foci in a transconjugant cell after acquisition of the F *ssb$^F$-sfgfp* plasmid. Scale bars 1 µm. Transconjugant (LY318 after acquisition of F *ssb$^F$-sfgfp* from LY270). (**E**) Histograms presenting the proportion of transconjugant cells with Ssb$^F$-sfGFP foci at the moment of mCh-ParB appearance, first and second duplication events. The mean and SD calculated from the indicated number (n) of transconjugant cells from at least three independent experiments (black dots) are shown.

| Strain | Relevant genotype[a, b, c] | Source or reference[a] |
|---|---|---|
| **MG1655 and derivatives** | | |
| **LY110** | MS388 / F-Tn*10*, *parS$_{PMT1}$-FRT-cat-FRT* | [1] |
| **LY117** | MS388 *ssb-ypet-FRT-kan-FRT* | [1] |
| **LY128** | MS388 *ssb-ypet-FRT* | [1] |
| **LY159** | MS388 / F-Tn*10*, *parS$_{PMT1}$-FRT*, Δ*ygeA::FRT-kan-FRT* | Conjugation LY864 x MS388 to St$^R$ Tc$^r$ |
| **LY160** | MS388 / F-Tn*10*, *parS$_{PMT1}$-FRT*, Δ*ygeA::FRT* | Derivative of LY159, kan removed via pCP20 |
| **LY240** | MS388 / F-Tn*10*, *parS$_{PMT1}$-FRT-cat-FRT*, *ssb$^F$-sfgfp-FRT-kan-FRT* | Conjugation LY225 x MS388 to St$^R$ Tc$^r$ |
| **LY242** | MS388 / F-Tn*10*, *parS$_{PMT1}$-FRT-cat-FRT*, *traM-sfgfp-FRT-kan-FRT* | Conjugation LY227 x MS388 to St$^R$ Tc$^r$ |
| **LY260** | MS388 / F-Tn*10*, *parS$_{PMT1}$-FRT-cat-FRT*, *traS-sfgfp-FRT-kan-FRT* | Conjugation LY149 x MS388 to St$^R$ Tc$^r$ |
| **LY261** | MS388 / F-Tn*10*, *parS$_{PMT1}$-FRT-cat-FRT*, *traT-sfgfp-FRT-kan-FRT* | Conjugation LY148 x MS388 to St$^R$ Tc$^r$ |
| **LY270** | MS388 / F-Tn*10*, *parS$_{PMT1}$-FRT*, *ssb$^F$-sfgfp-FRT* | Derivative of LY240, cat and kan removed via pCP20 |
| **LY272** | MS388 / F-Tn*10*, *parS$_{PMT1}$-FRT*, *traM-sfgfp-FRT* | Derivative of LY242, cat and kan removed via pCP20 |
| **LY284** | MS388 / F-Tn*10*, *parS$_{PMT1}$-FRT-cat-FRT*, *sopB-sfgfp-FRT-kan-FRT* | Conjugation LY276 x MS388 to St$^R$ Tc$^r$ |
| **LY318** | MS388 / pSN70 | [1] |
| **LY341** | MS388 / F-Tn*10*, *parS$_{PMT1}$-FRT-cat-FRT*, *traC-sfgfp-FRT-kan-FRT* | Conjugation LY327 x MS388 to St$^R$ Tc$^r$ |
| **LY355** | MS388 *ssb-ypet-FRT*; *mcherry-dnaN-frt-Kan-frt* | Transduction of *mcherry-dnaN-frt-Kan-frt* into LY128 to Km$^r$ |
| **LY358** | MS388 *ssb-ypet-FRT* / pSN70 | [1] |
| **LY390** | MS388 / F-Tn*10*, *parS$_{PMT1}$-FRT-cat-FRT*, *repE::PlacIQ1-sfgfp-FRT-kan-FRT* | [1] |
| **LY728** | MS388 / F-Tn*10*, *parS$_{PMT1}$-FRT*, Δ*ssb$^F$::FRT-kan-FRT* | Conjugation LY718 x MS388 to St$^R$ Tc$^r$ |
| **LY735** | MS388 / F-Tn*10*, *parS$_{PMT1}$-FRT*, *yfjB-sfgfp-FRT-kan-FRT* | Conjugation LY731 x MS388 to St$^R$ Tc$^r$ |
| **LY736** | MS388 / F-Tn*10*, *parS$_{PMT1}$-FRT*, *yfjA-sfgfp-FRT-kan-FRT* | Conjugation LY733 x MS388 to St$^R$ Tc$^r$ |
| **LY737** | MS388 / F-Tn*10*, *parS$_{PMT1}$-FRT*, *ygeA-sfgfp-FRT-kan-FRT* | Conjugation LY732 x MS388 to St$^R$ Tc$^r$ |
| **LY738** | MS388 / F-Tn*10*, *parS$_{PMT1}$-FRT*, *psiB-sfgfp-FRT-kan-FRT* | Conjugation LY734 x MS388 to St$^R$ Tc$^r$ |
| **LY754** | MS388 / F-Tn*10*, *parS$_{PMT1}$-FRT*, *psiB-sfgfp-FRT* | Derivative of LY738, kan removed via pCP20 |
| **LY755** | MS388 / F-Tn*10*, *parS$_{PMT1}$-FRT*, Δ*ssb$^F$::FRT* | Derivative of LY728, kan removed via pCP20 |
| **LY756** | MS388 / F-Tn*10*, *parS$_{PMT1}$-FRT*, *ygeA-sfgfp-FRT* | Derivative of LY737, kan removed via pCP20 |
| **LY791** | MS388 / F-Tn*10*, *parS$_{PMT1}$-FRT*, *yfjA-sfgfp-FRT* | Derivative of LY736, kan removed via pCP20 |
| **LY792** | MS388 / F-Tn*10*, *parS$_{PMT1}$-FRT*, *yfjB-sfgfp-FRT* | Derivative of LY735, kan removed via pCP20 |

| | | |
|---|---|---|
| **LY803** | MS388 / F-Tn*10*, *parS$_{PMT1}$-FRT, ΔFrpo1::FRT-kan-FRT* | Conjugation LY795 x MS388 to St$^R$ Tc$^r$ |
| **LY804** | MS388 / F-Tn*10*, *parS$_{PMT1}$-FRT, ΔFrpo2::FRT-kan-FRT* | Conjugation LY796 x MS388 to St$^R$ Tc$^r$ |
| **LY823** | MS388 / F-Tn*10*, *parS$_{PMT1}$-FRT, ΔFrpo2::FRT* | Derivative of LY804, kan removed via pCP20 |
| **LY824** | MS388 / F-Tn*10*, *parS$_{PMT1}$-FRT, ΔFrpo1::FRT* | Derivative of LY803, kan removed via pCP20 |
| **LY834** | MS388 / F-Tn*10*, *parS$_{PMT1}$-FRT*, *sopB-sfgfp-FRT* | Derivative of LY284, cat and kan removed via pCP20 |
| **LY875** | MS388 / F-Tn*10*, *parS$_{PMT1}$-FRT*, | Conjugation LY162 x MS388 to St$^R$ Tc$^r$ |
| **LY1007** | MS388 *ssb-ypet-FRT* / F-Tn*10*, *parS$_{PMT1}$-FRT* | Conjugation LY162 x LY128 to St$^R$ Tc$^r$ |
| **LY1038** | MS388 / F-Tn*10*, *parS$_{PMT1}$-FRT*, *ssb$^F$-mcherry-FRT-kan-FRT* | Conjugation LY1037 x MS388 to St$^R$ Tc$^r$ |
| **LY1053** | MS388 / F-Tn*10*, *parS$_{PMT1}$-FRT*, *ssb$^F$-mcherry-FRT* | Derivative of LY038, kan removed via pCP20 |
| **LY1068** | MS388 *ssb-ypet-FRT* / F-Tn*10*, *parS$_{PMT1}$-FRT, Δssb$^F$::FRT* | Conjugation LY837 x LY128 to St$^R$ Tc$^r$ |
| **LY1149** | MS388 / F-Tn*10*, *parS$_{PMT1}$-FRT*, *traC-sfgfp-FRT* | Derivative of LY341, cat and kan removed via pCP20 |
| **LY1150** | MS388 / F-Tn*10*, *parS$_{PMT1}$-FRT*, *traS-sfgfp-FRT* | Derivative of LY260, cat and kan removed via pCP20 |
| **LY1151** | MS388 / F-Tn*10*, *parS$_{PMT1}$-FRT*, *traT-sfgfp-FRT* | Derivative of LY261, cat and kan removed via pCP20 |
| **LY1216** | MS388 / F-Tn*10*, *parS$_{PMT1}$-FRT*, *repE::P$_{lacIQ1}$-sfgfp-FRT* | Derivative of LY390, cat and kan removed via pCP20 |
| **LY1224** | MS388 / F-Tn*10*, *parS$_{PMT1}$-FRT*, *ygfA-sfgfp-FRT-kan-FRT* | Conjugation LY1219 x MS388 to St$^R$ Tc$^r$ |
| **LY1225** | MS388 / F-Tn*10*, *parS$_{PMT1}$-FRT*, *traM::P$_{lacIQ1}$-sfgfp-FRT-kan-FRT* | Conjugation LY1220 x MS388 to St$^R$ Tc$^r$ |
| **LY1226** | MS388 / F-Tn*10*, *parS$_{PMT1}$-FRT*, *tnpA::PlacIQ1-sfgfp-FRT-kan-FRT* | Conjugation LY1221 x MS388 to St$^R$ Tc$^r$ |
| **LY1230** | MS388 / F-Tn*10*, *parS$_{PMT1}$-FRT*, *ygfA-sfgfp-FRT* | Derivative of LY1224, kan removed via pCP20 |
| **LY1231** | MS388 / F-Tn*10*, *parS$_{PMT1}$-FRT*, *traM::P$_{lacIQ1}$-sfgfp-FRT* | Derivative of LY1225, kan removed via pCP20 |
| **LY1232** | MS388 / F-Tn*10*, *parS$_{PMT1}$-FRT*, *tnpA::P$_{lacIQ1}$-sfgfp-FRT* | Derivative of LY1226, kan removed via pCP20 |
| **LY1288** | MS388 *rpsL* (St$^R$), *ssb-ypet-FRT*, *dnaE-kan* | P1 LY1158 x LY128 to Kan$^R$ ts |
| **LY1359** | MS388 *rpsL* (St$^R$), *ssb-ypet-FRT*, *polA-tc* | P1 LY1159 x LY128 to Tc$^R$ ts |
| **LY1364** | MS388 / F-Tn*10*, *parS$_{PMT1}$-FRT*, *ΔFrpo2::FRT, yfjA-sfgfp-FRT-kan-FRT* | Conjugation LY1355 x MS388 to St$^R$ Tc$^r$ |
| **LY1365** | MS388 / F-Tn*10*, *parS$_{PMT1}$-FRT*, *ΔFrpo2::FRT, ssb$^F$-sfgfp-FRT-kan-FRT* | Conjugation LY1356 x MS388 to St$^R$ Tc$^r$ |
| **LY1368** | MS388 / F-Tn*10*, *parS$_{PMT1}$-FRT*, *ΔFrpo1::FRT, ygeA-sfgfp-FRT-kan-FRT* | Conjugation LY1367 x MS388 to St$^R$ Tc$^r$ |
| **LY1373** | MS388 *rpsL* (St$^R$), *ssb-ypet-FRT* / pSN70, clone 1 | pSN70 x LY1288 to Ap$^r$ |

| | | |
|---|---|---|
| **LY1374** | MS388 *rpsL* (St$^R$), *ssb-ypet-FRT* / pSN70, clone 2 | pSN70 x LY1288 to Ap$^r$ |
| **LY1382** | MS388 *rpsL* (St$^R$), *ssb-ypet-FRT /* pAC1 | pAC1 x LY128 to Ap$^r$ |
| **LY1418** | MS388 / F-Tn*10*, *parS$_{PMT1}$-FRT*, Δ*Frpo1-ygeA::FRT-kan-FRT* | Conjugation LY1414 x MS388 to St$^R$ Tc$^r$ |
| **LY1419** | MS388 / F-Tn*10*, *parS$_{PMT1}$-FRT*, Δ*Frpo1-ygeA-ygeB::FRT-kan-FRT* | Conjugation LY1415 x MS388 to St$^R$ Tc$^r$ |
| **LY1424** | MS388 / F-Tn*10*, *parS$_{PMT1}$-FRT*, Δ*Frpo1-ygeA::FRT* | Derivative of LY1418, kan removed via pCP20 |
| **LY1425** | MS388 / F-Tn*10*, *parS$_{PMT1}$-FRT*, Δ*Frpo1-ygeA-ygeB::FRT* | Derivative of LY1419, kan removed via pCP20 |
| **MG1655** | $\lambda^-$, *rph-1* | Coli Genetic Stock Center (CGSC) #6300 |
| **MS388** | MG1655 *rpsL* (St$^R$) | Gift from F. Cornet |
| **MS428** | MG1655 *rpsL* (St$^R$), ΔlacZ | Gift from F. Cornet |
| **Other genetic backgrounds** | | |
| **K603** | F$^+$ [F1-10(Tn*10*)], *thr-1, araC14, leuB6*(Am)*, lacY1, glnX44*(AS)*, galK2*(Oc)*, galT22, $\lambda^-$, ΔtrpE63, xylA5, mtl-1, thiE1* | Coli Genetic Stock Center CGSC #6451 Strain carrying the F-Tn*10* natural isolate with Tn*10* transposon in the intergenic region *ybdB-ybfA*. GenBank accession number MK492260) |
| **DY330** | W3110 Δ*lacU169, gal490, λcI857,* Δ*(cro-bioA)* | [2] |
| **LY183** | AB1157 kan-11aa-mcherry-dnaN = | [3] |
| **LY162** | DY330 / F-Tn*10-parS$_{PMT1}$-FRT* | [1] |
| **LY148** | DY330 / F-Tn*10*, *parS$_{PMT1}$-FRT-cat-FRT*, *traT-sfgfp-FRT-kan-FRT* | λred *traT-sfgfp* fusion at the endogenous F plasmid *locus* (OL136/OL137) |
| **LY149** | DY330 / F-Tn*10*, *parS$_{PMT1}$-FRT-cat-FRT*, *traS-sfgfp-FRT-kan-FRT* | λred *traS-sfgfp* fusion at the endogenous F plasmid *locus* (OL138/OL139) |
| **LY225** | DY330 / F-Tn*10*, *parS$_{PMT1}$-FRT-cat-FRT*, *ssb$^F$-sfgfp-FRT-kan-FRT* | λred *ssb$^F$-sfgfp* fusion at the endogenous F plasmid *locus* (OL25/OL26) |
| **LY227** | DY330 / F-Tn*10*, *parS$_{PMT1}$-FRT-cat-FRT*, *traM-sfgfp-FRT-kan-FRT* | λred *traM-sfgfp* fusion at the endogenous F plasmid *locus* (OL21/OL22) |
| **LY276** | DY330 / F-Tn*10*, *parS$_{PMT1}$-FRT-cat-FRT*, *sopB-sfgfp-FRT-kan-FRT* | λred *sopB-sfgfp* fusion at the endogenous F plasmid *locus* (OL17/OL18) |
| **LY327** | DY330 / F-Tn*10*, *parS$_{PMT1}$-FRT-cat-FRT*, *traC-sfgfp-FRT-kan-FRT* | λred *traC-sfgfp* fusion at the endogenous F plasmid *locus* (OL189/OL190) |
| **LY404** | DY330 / F-Tn*10*, *parS$_{PMT1}$-FRT-cat-FRT*, *repE::P$_{lacIQ1}$-sfgfp-FRT-kan-FRT* | [1] |
| **LY718** | DY330 / F-Tn*10*, *parS$_{PMT1}$-FRT*, Δ*ssb$^F$::FRT-kan-FRT* | λred Δ*ssb$^F$::kan* construct at the endogenous F plasmid *locus* (OL194/OL195) |
| **LY731** | DY330 / F-Tn*10*, *parS$_{PMT1}$-FRT*, *yfjB-sfgfp-FRT-kan-FRT* | λred *yfjB-sfgfp* fusion at the endogenous F plasmid *locus* (OL418/OL419) |
| **LY732** | DY330 / F-Tn*10*, *parS$_{PMT1}$-FRT*, *ygeA-sfgfp-FRT-kan-FRT* | λred *ygeA-sfgfp* fusion at the endogenous F plasmid *locus* (OL422/OL423) |
| **LY733** | DY330 / F-Tn*10*, *parS$_{PMT1}$-FRT*, *yfjA-sfgfp-FRT-kan-FRT* | λred *yfjA-sfgfp* fusion at the endogenous F plasmid *locus* (OL415/OL416) |
| **LY734** | DY330 / F-Tn*10*, *parS$_{PMT1}$-FRT*, *psiB-sfgfp-FRT-kan-FRT* | λred *psiB-sfgfp* fusion at the endogenous F plasmid *locus* (OL429/OL430) |
| **LY795** | DY330 / F-Tn*10*, *parS$_{PMT1}$-FRT*, Δ*Frpo1::FRT-kan-FRT* | λred Δ*Frpo1 :: kan* construct at the endogenous F plasmid *locus* (OL476/OL477) |

| | | |
|---|---|---|
| **LY796** | DY330 / F-Tn*10*, *parS$_{PMT1}$-FRT*, Δ*Frpo2::FRT-kan-FRT* | λred Δ*Frpo2 :: kan* construct at the endogenous F plasmid *locus* (OL473/OL474) |
| **LY837** | DY330 / F-Tn*10*, *parS$_{PMT1}$-FRT*, Δ*ssB$^F$::FRT* | Conjugation LY755 x DY330 to $St^s$ $Tc^r$ |
| **LY864** | DY330 / F-Tn*10*, *parS$_{PMT1}$-FRT*, Δ*ygeA::FRT-kan-FRT* | λred Δ*ygeA :: kan* construct at the endogenous F plasmid *locus* (OL476/OL508) |
| **LY916** | DY330 / F-Tn*10*, *parS$_{PMT1}$-FRT,* Δ*Frpo1::FRT* | Conjugation LY824 x DY330 to $St^s$ $Tc^r$ |
| **LY917** | DY330 / F-Tn*10*, *parS$_{PMT1}$-FRT,* Δ*Frpo2::FRT* | Conjugation LY823 x DY330 to $St^s$ $Tc^r$ |
| **LY1037** | DY330 / F-Tn*10*, *parS$_{PMT1}$-FRT*, *ssb$^F$-mcherry-FRT-kan-FRT* | λred *ssb$^F$-mcherry* fusion at the endogenous F plasmid *locus* (OL92/OL28) |
| **LY1219** | DY330 / F-Tn*10*, *parS$_{PMT1}$-FRT*, *ygfA-sfgfp-FRT-kan-FRT* | λred *ygfA-sfgfp* fusion at the endogenous F plasmid *locus* (OL621/OL622) |
| **LY1220** | DY330 / F-Tn*10*, *parS$_{PMT1}$-FRT*, *traM::P$_{lacIQ1}$-sfgfp-FRT-kan-FRT* | λred *P$_{lacIQ1}$-sfgfp-FRT-kan-FRT* construct at the F plasmid intergenic *traM locus* (OL114/OL115) |
| **LY1221** | DY330 / F-Tn*10*, *parS$_{PMT1}$-FRT*, *tnpA::P$_{lacIQ1}$-sfgfp-FRT-kan-FRT* | λred *P$_{lacIQ1}$-sfgfp-FRT-kan-FRT* construct at the F plasmid intergenic *traM locus* (OL623/OL624) |
| **LY1355** | DY330 / F-Tn*10*, *parS$_{PMT1}$-FRT*, Δ*Frpo2::FRT, yfjA-sfgfp-FRT-kan-FRT* | λred *yfjA-sfgfp* fusion at the endogenous F plasmid *locus* of LY917 (OL415/OL416) |
| **LY1356** | DY330 / F-Tn*10*, *parS$_{PMT1}$-FRT*, Δ*Frpo2::FRT, ssb$^F$-sfgfp-FRT-kan-FRT* | λred *ssb$^F$-sfgfp* fusion at the endogenous F plasmid *locus* of LY917 (OL25/OL26) |
| **LY1367** | DY330 / F-Tn*10*, *parS$_{PMT1}$-FRT*, Δ*Frpo1::FRT, ygeA-sfgfp-FRT-kan-FRT* | λred *ygeA-sfgfp* fusion at the endogenous F plasmid *locus* of LY916 (OL422/OL423) |
| **LY1414** | DY330 / F-Tn*10*, *parS$_{PMT1}$-FRT*, Δ*Frpo1-ygeA::FRT-kan-FRT* | λred Δ*Frpo1-ygeA :: kan* construct at the endogenous F plasmid *locus* (OL507/OL508) |
| **LY1415** | DY330 / F-Tn*10*, *parS$_{PMT1}$-FRT*, Δ*Frpo1-ygeA-ygeB::FRT-kan-FRT* | λred Δ*Frpo1-ygeA-ygeB :: kan* construct at the endogenous F plasmid *locus* (OL476/OL514) |

[a] The abbreviations *bla*, *kan and cat* refer to insertions genes conferring resistance to ampicillin ($Ap^r$) kanamycin ($Kn^r$), and chloramphenicol ($Cm^r$); *rpsl* refers to spontaneous mutation conferring resistance to streptomycin. *FRT* refers to the FLP site-specific recombination site.

[b] Tn*10* transposon is located in the intergenic region *ybdB-ybfA* on the F plasmid.

[c] *sfgfp* gene encodes the superfolder Green Fluorescent Protein sfGFP

**Table S1. Strains list**

| Name | Construct and Usage [a] | Source or reference |
|---|---|---|
| **pCP20** | Flp expression plasmid | [4] |
| **p mCherry-*ParB* (pSN70)** | IPTG inducible expression of N-terminal fusion mCherry-$ParB_{PMT1}$ | [1] |
| **pR6K-sfGFP** | Carries *sfgfp-FRT-kan-FRT* used for several C-terminal fusion by λred | [1] |
| **pROD62** | Carries *mCherry-FRT-kan-FRT* used for Δ*Frpo1:: kan* and Δ*Frpo2 :: kan* deletion into F-Tn*10*, *parS*$_{PMT1}$*-FRT* (OL473/OL474, OL476/OL477) by λred and for plasmids construction by Gibson assembly | Gift from R. Reyes-Lamothe, McGill University |
| **pAC1** | Carries *ssb*$^{F}$*-mCherry* insertion under $P_{trc}$ promotor on pTrc99a by Gibson assembly (OL608/OL653, OL602/OL603) | Fig. S7 |
| **F-Tn*10* conjugative plasmid (from K603) derivatives** | | |
| **F*wt*** | F-Tn*10* with *parS*$_{PMT1}$ inserted at the intergenic *ygeB-ygfA locus* | [1] Fig. 1-2, 4B, 5B and Fig. S1-3 |
| **F *ssb*$^{F}$*-sfgfp*** | F-Tn*10* *parS*$_{PMT1}$ with *ssb*$^{F}$*-sfgfp* translational fusion at the endogenous locus | Fig. 3-4 and Fig. S4-5 |
| **F *ygeA-sfgfp*** | F-Tn*10* *parS*$_{PMT1}$ with *ygeA-sfgfp* translational fusion at the endogenous locus | Fig. 3-4 and Fig. S4-5 |
| **F *yfjA-sfgfp*** | F-Tn*10* *parS*$_{PMT1}$ with *yfjA-sfgfp* translational fusion at the endogenous locus | Fig. 3-4 and Fig. S4-5 |
| **F *yfjB-sfgfp*** | F-Tn*10* *parS*$_{PMT1}$ with *yfjB-sfgfp* translational fusion at the endogenous locus | Fig. 3 and Fig. S4-5 |
| **F *psiB-sfgfp*** | F-Tn*10* *parS*$_{PMT1}$ with *psiB-sfgfp* translational fusion at the endogenous locus | Fig. 3 and Fig. S4-5 |
| **F *sopB-sfgfp*** | F-Tn*10* *parS*$_{PMT1}$ with *sopB-sfgfp* translational fusion at the endogenous locus | Fig. 3 and Fig. S4-5 |
| **F *ygfA-sfgfp*** | F-Tn*10* *parS*$_{PMT1}$ with *ygfA-sfgfp* translational fusion at the endogenous locus | Fig. 3 and Fig. S4-5 |
| **F *traM-sfgfp*** | F-Tn*10* *parS*$_{PMT1}$ with *traM-sfgfp* translational fusion at the endogenous locus | Fig. 3 and Fig. S4-5 |
| **F *traT-sfgfp*** | F-Tn*10* *parS*$_{PMT1}$ with *traT-sfgfp* translational fusion at the endogenous locus | Fig. 3 and Fig. S4-5 |
| **F *traC-sfgfp*** | F-Tn*10* *parS*$_{PMT1}$ with *traC-sfgfp* translational fusion at the endogenous locus | Fig. 3 and Fig. S4-5 |
| **F *traS-sfgfp*** | F-Tn*10* *parS*$_{PMT1}$ with *traS-sfgfp* translational fusion at the endogenous locus | Fig. 3 and Fig. S4-5 |

| | | |
|---|---|---|
| **F *repE*::P$_{lacIQ1}$-sfGFP** | F-Tn*10 parS*$_{PMT1}$ with insertion *P*$_{lacIQ1}$-*sfgfp* in the *repE-sopA* intergenic region | Fig. 3 and Fig. S5 |
| **F *traM*::P$_{lacIQ1}$-sfGFP** | F-Tn*10 parS*$_{PMT1}$ with insertion *P*$_{lacIQ1}$-*sfgfp* in the *traM-traJ* intergenic region | Fig. 3 and Fig. S5 |
| **F *tnpA*::P$_{lacIQ1}$-sfGFP** | F-Tn*10 parS*$_{PMT1}$ with insertion *P*$_{lacIQ1}$-*sfgfp* in the *tnpA-ybaA* intergenic region | Fig. 3 and Fig. S5 |
| **F Δ*Frpo2*** | F-Tn*10 parS*$_{PMT1}$ with *Frpo2* deletion | Fig. 4B |
| **F Δ*Frpo1*** | F-Tn*10 parS*$_{PMT1}$ with *Frpo1* deletion | Fig. 4B |
| **F Δ*Frpo2 yfjA-sfgfp*** | F-Tn*10 parS*$_{PMT1}$ with *yfjA-sfgfp* translational fusion at the endogenous locus and *Frpo2* promotor deletion | Fig. 4 and Fig. S6B |
| **F Δ*Frpo2 ssb*$^{F}$*-sfgfp*** | F-Tn*10 parS*$_{PMT1}$ with *ssb*$^{F}$*-sfgfp* translational fusion at the endogenous locus and *Frpo2* promotor deletion | Fig. 4 and Fig. S6B |
| **F Δ*Frpo1 ygeA-sfgfp*** | F-Tn*10 parS*$_{PMT1}$ with *ygeA-sfgfp* translational fusion at the endogenous locus and *Frpo1* promotor deletion | Fig. 4 and Fig. S6B |
| **F Δ*ssb*$^{F}$** | F-Tn*10 parS*$_{PMT1}$ with *ssb*$^{F}$ *deletion* | Fig. 4 |
| **F Δ*Frpo1* Δ*ygeA*** | F-Tn*10 parS*$_{PMT1}$ with *Frpo1* and *ygeA* deletions | Fig. 4 |
| **F Δ*Frpo1* Δ*ygeAB*** | F-Tn*10 parS*$_{PMT1}$ with *Frpo1, ygeA* and *ygeB* deletions | Fig. 4 |

[a] *sfgfp* gene encodes the superfolder Green Fluorescent Protein sfGFP

**Table S2. Plasmids used in this study**

| Name | Sequence | Construct |
|---|---|---|
| **OL1** | GTATTGTCACTCATATGATATTTTTGTGTGTG<br>GCCTTCCAGGTCTGCTATGTGGTGCTATCT | λred *parS*$_{PMT1}$*-FRT-cat-FRT* insertion at the F plasmid intergenic *ygeB-ygfA locus*. PCR on pGBKD3-parS |
| **OL2** | ACAGGCATTGTCAGATACCGGTTATGCCGCA<br>AAAGCGGCAGATTGTGTAGGCTGGAGCTGC | |
| **OL17** | ATTGAGGCCATTCTTAAGGAACTTGAAAAGC<br>CAGCACCCAGCTCGGCTGGCTCCGCT | λred *sopB-sfgfp* and *sopB-mTurquoise* fusion at the endogenous F plasmid *locus*. PCR on pR6K-sfGFP and pr6K-mTurquoise plasmids. |
| **OL18** | TAAGTAAAGACAGATAAACGTAGACTAAAA<br>CGTGGTCGCACATATGAATATCCTCCTTAG | |
| **OL21** | ATCTGAGATGGAACGATTTTTCCAAAAAAT<br>GATGATGAAAGCTCGGCTGGCTCCGCT | λred *traM-sfgfp* fusion at the endogenous F plasmid *locus*. PCR on pR6K-sfGFP plasmid. |
| **OL22** | ATCATAAAAACTCTGATATTTGAACGAAGTC<br>AAATTTCGTCATATGAATATCCTCCTTAG | |
| **OL25** | GGAGGGTGACGATTACGGGTTTTCAGACGAT<br>ATCCCGTTCAGCTCGGCTGGCTCCGCT | λred *ssb*$^{F}$*-sfgfp* fusion at the endogenous F plasmid *locus*. PCR on pR6K-sfGFP plasmid. |
| **OL26** | TGCCCCGCACAGGACGGGGCGGTTGTCACAG<br>TCAGTCGTCATATGAATATCCTCCTTAG | |
| **OL36** | TGGTTCTGGCGAATTCGTGTCTAAAGGTGAA<br>GAACTGTTCACCG | pR6K-sfGFP plasmid construct using Gibson assembly. PCR on pROD62 (OL38/OL39) and pKD4-sfGFP (OL36/OL37) |
| **OL37** | GCTCCAGCCTACACCCGGGTTATTTGTAGAG<br>CTCATCCAT | |
| **OL38** | CCCGGGTGTAGGCTGGAG | |
| **OL39** | CACGAATTCGCCAGAACCAGC | |
| **OL114** | CCAAAAAATGATGATGAATAAACGAAATTTG<br>ACTTCGTTCCGACTGGAAAGCGGGCAGTG | λred $P_{lacIQ1}$*-sfgfp-FRT-kan-FRT* construct at F plasmid intergenic *traM locus*. PCR on pR6K-sfGFP plasmid |
| **OL115** | CGTACTGTCACCTTTTTAAATCATAAAAACTC<br>TGATATTTCATATGAATATCCTCCTTAG | |
| **OL116** | ATTTCTTCTTGCGCTGAGCGTAAGAGCTATCT<br>GACAGAACCGACTGGAAAGCGGGCAGTG | λred $P_{lacIQ1}$*-sfgfp-FRT-kan-FRT* construct at F plasmid intergenic *repE-sopA locus*. PCR on pR6K-sfGFP plasmid |
| **OL117** | GAGCATAGCGAGCGAACTGGCGAGGAAGCA<br>AAGAAGAACTCATATGAATATCCTCCTTAG | |
| **OL136** | TCTCGAAGACCAGCTGGCCAAATCAATCGCA<br>AATATTCTCAGCTCGGCTGGCTCCGC T | λred *traT-sfgfp* fusion at the endogenous F plasmid *locus*. PCR on pR6K-sfGFP plasmid |
| **OL137** | GTCAGTCAGGAGGCCGGTCAGACCAGCCTCC<br>GGAAGATAACATATGAATATCCTCCTTAG | |
| **OL138** | CAATAGTTACGTCAAAAACAAGAAGTTATCA<br>AGAGTAAAAAGCTCGGCTGGCTCCGCT | λred *traS-sfgfp* fusion at the endogenous F plasmid *locus*. PCR on pR6K-sfGFP plasmid |
| **OL139** | GTTTTTTTGTTTTCATCATATATTTACTCTCTA<br>ATATCTTCATATGAATATCCTCCTTAG | |
| **OL189** | AGCCTGGCTGGAAGAACATGAGAAATACAG<br>GAGTGTGGCAAGCTCGGCTGGCTCCGCT | λred *traC-sfgfp* fusion at the endogenous F plasmid *locus*. PCR on pR6K-sfGFP plasmid |
| **OL190** | GTTCTGCCGTGACGTCGGCGGGTTTCTGCGTT<br>GAACTCATCATATGAATATCCTCCTTAG | |
| **OL194** | ATGGGGGCTGAAAACATCAACAGAAGGAGA<br>CACATCGTGTAGGCTGGAGCTGCTTC | λred Δ*ssb*$^{F}$*::kan* construct at the endogenous F plasmid *locus*. PCR on pROD62 |
| **OL195** | ATGCCCCGCACAGGACGGGGCGGTTGTCACA<br>GTCAGTCGTCATATGAATATCCTCCTTAG | |
| **OL415** | CTGGATGAACCATGTCATCCGTGATTTCGGG<br>ATTCTTAAGAGCTCGGCTGGCTCCGCTGC | λred *yfjA-sfgfp* fusion at the endogenous F plasmid *locus*. PCR on pR6K-sfGFP plasmid |
| **OL416** | AATGATTGTCTGCCGGACGCGTTCCCGTCCG<br>GCACCTCTCTATCCTCCTTAGTTCCTATT | |

| | | |
|---|---|---|
| **OL418** | GGATGCCACTGAACGTACCGATAACCTGGCTGATGCCGCCAGCTCGGCTGGCTCCGCTGC | $\lambda$red *yfjB-sfgfp* fusion at the endogenous F plasmid *locus*. PCR on pR6K-sfGFP plasmid |
| **OL419** | TGCTGCCGCCCCGTCTCCGGCGGGGCGGTGTGGTTGTTCATATCCTCCTTAGTTCCTATT | |
| **OL422** | TCAGATTAAGCGTGCCGGCCTCGATATCCGGGCGATCTGCAGCTCGGCTGGCTCCGCTGC | $\lambda$red *ygeA-sfgfp* fusion at the endogenous F plasmid *locus*. PCR on pR6K-sfGFP plasmid |
| **OL423** | CAAGATTGCAACAATCAGGAGGGATATTCATCACATCCGGTATCCTCCTTAGTTCCTATT | |
| **OL429** | CGACATCATCCATATCCTGATGGCGGAAGGAGGTCAGGTAAGCTCGGCTGGCTCCGCTGC | $\lambda$red *psiB-sfgfp* fusion at the endogenous F plasmid *locus*. PCR on pR6K-sfGFP plasmid |
| **OL430** | CTGTGCTGAGGGGAACCAGTGCCTGTGAACGTACGCTCATTATCCTCCTTAGTTCCTATT | |
| **OL473** | CTTCCTCTTCTTCCTCTTCTTTCCGTTCCTCTCCTGCTTAGTGTAGGCTGGAGCTGCTTC | $\lambda$red $\Delta$*Frpo2::kan* construct at the endogenous F plasmid *locus*. PCR on pROD62 |
| **OL474** | TAATGCCACGAACTGCCATGATGTGTCTCCTTCTGTTGATCATATGAATATCCTCCTTAG | |
| **OL476** | GCTGCCCTGATTCCTGTCTGTTTTGTTGTTCTCCTGGCAGGTGTAGGCTGGAGCTGCTTC | $\lambda$red $\Delta$*Frpo1::kan* construct at the endogenous F plasmid *locus*. PCR on pROD62 |
| **OL477** | TGGATATTTCGGGTACTCATCCTGCTTTCTCCTCTCTCTGCATATGAATATCCTCCTTAG | |
| **OL507** | AGATGATGGGGGCTGAAAACCAGAGAGAGGAGAAAGCAGGGTGTAGGCTGGAGCTGCTTC | $\lambda$red $\Delta$*ygeA::kan* construct at the endogenous F plasmid *locus*. PCR on pROD62 |
| **OL508** | CAAGATTGCAACAATCAGGAGGGATATTCATCACATCCGGCATATGAATATCCTCCTTAG | |
| **OL514** | TGATAATATAAGTAAAGCCCCGAAATTTTTTCGGGGCTTTACTTATATTACATATGAATATCCTCCTTAG | $\lambda$red $\Delta$*FrpO-ygeA-ygeB::kan* construct at the endogenous F plasmid *locus*. PCR on pROD62 |
| **OL602** | GATCCTCTAGAGTCGACCTGCAGGC | pAC1 plasmid construct using Gibson assembly. PCR on pTrc99a (OL602/OL603) and on LY1053 (OL608/OL653) |
| **OL603** | CATGGTCTGTTTCCTGTGTGAAATT | |
| **OL608** | AACAATTTCACACAGGAAACAGACCATGGCAGTTCGTGGCATTAA | |
| **OL653** | GCCTGCAGGTCGACTCTAGAGGATCTTACTTGTACAGCTCGTCCA | |
| **OL621** | GAAGAAATCACTTCCGGAAATTAACAGCGTGCAGAACAATAGCTCGGCTGGCTCCGCT | $\lambda$red *ygfA-sfgfp* fusion at the endogenous F plasmid *locus*. PCR on pR6K-sfGFP plasmid |
| **OL622** | GCTTTTGCGGCATAACCGGTATCTGACAATGCCTGTAAGACATATGAATATCCTCCTTAG | |
| **OL623** | GGGGGTGACTTGAAGTAAGCCTTATTTTTCTCCTCCTTTGCGACTGGAAAGCGGGCAGTG | $\lambda$red $P_{lacIQ1}$-*sfgfp-FRT-kan-FRT* construct at F plasmid intergenic *tnpA locus*. PCR on pR6K-sfGFP plasmid |
| **OL624** | GTTTATTGGTATGAAATAAGAAAAAAACCTCCTTTGAATTCATATGAATATCCTCCTTAG | |

**Table S3. PCR primers used for strain and plasmid constructions**

**Movie S1.**

Microfluidic time-lapse imaging of conjugation showing the production of leading proteins sfGFP fusion as respect to the formation of a mCh-ParB focus in transconjugant cells. Donors carry the F derivative with the indicated sfGFP fusion, and recipient cells produce mCh-ParB from the pSN70 plasmid. Before plasmid transfer, the mCh-ParB is diffuse in the recipient cells and mCh-ParB focus formation reports the ssDNA-to-dsDNA conversion after ssDNA acquisition. Merge of phase contrast, mCherry and sfGFP channels are shown. Cells were grown in M9-CAA at 37°C, images were taken every 5 min. Scale bar 1 µm and time in minutes are indicated.

**Movie S2.**

Microfluidic time-lapse imaging of conjugation showing the production of maintenance and Tra proteins sfGFP fusions as respect to the formation of a mCh-ParB focus in transconjugant cells. Donors carry the F derivative with the indicated sfGFP fusion, and recipient cells produce mCh-ParB from the pSN70 plasmid. Before plasmid transfer, the mCh-ParB is diffuse in the recipient cells and mCh-ParB focus formation reports the ssDNA-to-dsDNA conversion after ssDNA acquisition. Merge of phase contrast, mCherry and sfGFP channels are shown. Cells were grown in M9-CAA at 37°C, images were taken every 5 min. Scale bar 1 µm and time in minutes are indicated.

**Movie S3.**

Microfluidic time-lapse imaging of conjugation showing the production $Ssb^F$-sfGFP fusion as respect to the formation of a mCh-ParB focus in transconjugant cells. Donors carry the F *ssb$^F$-sfgfp* but do not exhibit green fluorescence as $Ssb^F$-sfGFP is not produced. $Ssb^F$-sfGFP is initially diffuse in the recipient cells when the mCh-ParB focus is already present. $Ssb^F$-sfGFP forms foci concomitantly with mCh-ParB focus duplication events. From left to right, merge of phase contrast, mCherry and sfGFP channels; merge of mCherry and sfGFP channels; mCherry channel; and sfGFP channel are shown. Cells were grown in M9-CAA at 37°C, images were taken every 5 min. Scale bar 1 µm and time in minutes are indicated.
